# HCF-2 inhibits cell proliferation and activates differentiation-gene expression programs

**DOI:** 10.1101/577890

**Authors:** Daria Gudkova, Oleksandr Dergai, Viviane Praz, Winship Herr

## Abstract

HCF-2 is a member of the <u>h</u>ost-<u>c</u>ell-<u>f</u>actor protein family, which arose in early vertebrate evolution as a result of gene duplication. Whereas its paralog, HCF-1, is known to act as a versatile chromatin-associated protein required for cell proliferation and differentiation, much less is known about HCF-2. Here, we show that HCF-2 is broadly present in human and mouse cells, and possesses activities distinct from HCF-1. Unlike HCF-1, which is excluded from nucleoli, HCF-2 is nucleolar — an activity conferred by one and a half C-terminal Fibronectin type 3 repeats and inhibited by the HCF-1 nuclear localization signal. Elevated HCF-2 synthesis in HEK-293 cells results in phenotypes reminiscent of HCF-1-depleted cells, including inhibition of cell proliferation and mitotic defects. Furthermore, increased HCF-2 levels in HEK-293 cells lead to inhibition of cell proliferation and metabolism gene-expression programs with parallel activation of differentiation and morphogenesis gene-expression programs. Thus, the HCF ancestor appears to have evolved into a small two-member protein family possessing contrasting nuclear vs. nucleolar localization, and cell proliferation and differentiation functions.

## INTRODUCTION

Gene duplication is a major evolutionary mechanism for creating genetic diversity (1). Such diversity is generated by subsequent mutation and divergence of the functions of each of the duplicated genes, in many cases resulting in gene families encoding proteins with opposing functions. Genes encoding transcription factors are prevalent among such duplicated-gene families (2, 3). For example, members of the E2F family, which play important roles in cell cycle control, differentiation and development (4), consist in mammals of both activator (e.g., E2F1, E2F2, and E2F3a) and repressor (e.g., E2F4 and E2F5) transcriptional regulators (5).

Here, we study HCF-1 and HCF-2, two proteins that resulted from gene duplication and in humans are encoded by the *HCFC1* and *HCFC2* genes. HCF-1, the more extensively studied of the two, acts as a <u>h</u>ost-<u>c</u>ell-<u>f</u>actor (HCF) protein for herpes simplex virus (HSV). It stabilizes formation of the so-called VP16-induced complex (VIC), which contains, besides HCF-1, the HSV virion protein VP16 and a second cellular transcriptional regulator called Oct-1 (reviewed by (6)). In uninfected cells, HCF-1 serves as a versatile transcriptional regulatory integrator, bringing together promoter-specific transcription factors with numerous chromatin modifiers facilitating either activation or repression of transcription (reviewed by 7). Human HCF-1 is synthesized as a large 2035-aa precursor protein, which then undergoes cleavage by *O*-linked-ß N-acetylglucosamine transferase (OGT) at any of six centrally located 20–26-aa repeats called HCF-1_PRO_ repeats (8–10). In mouse and human cells, HCF-1 controls the cell cycle at two points: (i) passage through the G1 phase, where it promotes cell cycle progression, and (ii) exit from mitosis, where it ensures proper chromosomal and nuclear segregation (11–13).

The lesser studied protein, HCF-2, is considerably smaller (human HCF-2 is 792 aa long) and yet shares important common features with HCF-1, namely N-terminal Kelch domain and C-terminal Fn3 repeats, which are highly conserved between the two proteins (14). Consistent with such sequence conservation, HCF-2 can also promote VIC formation, at least with a truncated VP16 protein lacking an acidic transcriptional activation domain (14). HCF-2 has also recently been implicated in transcriptional regulation as mutations in the mouse *Hcfc2* gene can abrogate HCF-2 involvement in interferon-regulatory-factor IRF-1 and IRF-2-dependent transcription (15). Thus, HCF-2 is an HCF-1 paralog that possesses shared but also novel activities. We probe these activities here and show that HCF-2 has acquired a prominent nucleolar localization as well as antiproliferative activities.

## MATERIALS AND METHODS

### Mammalian expression plasmids

Human *HCFC2* (*hHCFC2*) coding sequences from pCGN-HCF-2 (14) were subcloned into the vector pcDNA FRT/TO, with N-terminal Flag- and fluorescent Cherry-tag coding sequences, via ligation-independent cloning (16) to obtain pcDNA5FRT/TO/F-Cherry-HCF-2_FL_.

Mutations were created via QuickChange site-directed mutagenesis of pcDNA5FRT/TO/F-Cherry-HCF-2_FL_ (Agilent Technologies). Generation of the F-Cherry-HCF-2 Fn3c fragment and its derivatives was done by deletion and point mutagenesis. To obtain the HCF-1 Fn3c sequence, the HCF-1 Acidic region and NLS in pcDNA5/FRT-F-HCF-1_C600_ were deleted. Fn3n (W377A) and Fn3c (V653E) point mutations generated by site-directed mutagenesis in pcDNA5FRT/TO/F-Cherry-HCF-2_FL_ generated the F-Cherry-HCF-2_Fn3nc*_ coding sequences and insertion of the HCF-1 NLS sequence (2002-2035 aa) to the C-terminus of HCF-2 generated the F-Cherry-HCF-2_+NLS_ coding sequences. All constructs were verified by sequencing.

### Antibodies

Antibodies used in this study were (i) rabbit polyclonals α-HCF-1_C_ (H12) (17) and α-Cherry (183628, Abcam); (ii) mouse monoclonals α-HCF-1 (M2) (18), α-MEK2 (610236, BD Transduction Laboratories™), α-histone H3 (05-499, Upstate), anti-α-tubulin (32293, Santa Cruz Biotechnology), α-B23 (NPM) (56622, Santa Cruz Biotechnology), α-RPA194 (48385, Santa Cruz Biotechnology), α-Flag-M2 (1804, Sigma), and α-NCL (396400, Life Technologies); (iii) α-SC-35, rabbit monoclonal (204916, Abcam); and (iv) normal rabbit IgG (2027, Santa Cruz Biotechnology).

### HCF-2 antibody preparation

For anti-mouse HCF-2 (mHCF-2) antibody, the cDNA sequence encoding mHCF-2 aa 394-526 was amplified from mouse embryo fibroblast (MEF) cDNA with appropriate primers and inserted into the pET47b vector, using BamHI and NotI restriction sites, creating pET47b-6xHis-HCF-2_394-526_. In pET47b-6xHis-HCF-2_394-526_, the HCF-2_394-526_ segment is fused at its N-terminus to a 6xHis-tag followed by the human rhinovirus (HRV) protease recognition site.

His-tagged 6xHis-HCF-2_394-526_ protein was synthesized in pET47b-6xHis-HCF-2_394-526_-transformed BL21 (DE3) E. coli cells grown at 37°C by addition of 0.2 mM IPTG and native protein purified using Nickel affinity chromatography according to the manufacturer’s protocol (Qiagen). For N-terminal His-tag removal, Ni-NTA resin bound 6xHis-mHCF-2_394-526_ protein was treated with HRV 3C protease and the 6xHis tag left bound to the resin. After preparative PAGE and concentration with Amicon Ultra concentration tubes (Millipore), the protein was used for rabbit immunization by AbFrontier (South Korea).

### Immunoprecipitation and immunoblotting

Cell extracts were prepared by lysing cells in whole-cell-lysis (WCL) extraction buffer (10 mM Hepes, pH7.9, 250 mM NaCl, 0.25% NP-40, 5% glycerol, 0,2 mM EDTA, 50 uM NaF, 1 mM DTT) for 30 min at 4°C and further cleared by centrifugation at 13,000 rpm for 20 min at 4°C. For immunoprecipitation, 0.5-1 mg of cell extracts was incubated with 1-2 μg of indicated antibody for 3 h or overnight at 4°C followed by a 1 h incubation with protein A-sepharose beads. For immunobot analysis, samples were washed 3-4 times with extraction buffer, boiled in the 1 × Laemmli buffer and further analyzed by immunoblotting as described (8).

### HCF-2 LC-MS/MS analysis

For mass-spectroscopy (MS) analysis of immunoprecipitated HCF-2, 2 × 10^7^ MEF or 2 × 10^8^ human embryonic kidney-293 (HEK-293) cells were harvested and proteins extracted by treatment with WCL extraction buffer. HCF-2 proteins were immunoprecipitated by incubating the whole cell extract for 3 h with 2 ug α-HCF-2 antibody or normal rabbit IgG (as a negative control) followed by BSA-blocked agarose A beads for 1 h. The beads were washed 4 times with WCL buffer and boiled in 1 × Laemmli buffer. One tenth of the sample was used for analytical PAGE and the remainder purified by PAGE and the band corresponding to the predicted HCF-2 size (72 kDa for mHCF-2 and 100 kDa for hHCF-2) was cut out of the gel after Coomassie-staining and subjected to mass spectrometry after digestion with trypsin (19). For identification of proteins in HCF-2 complexes from MEF cells, 2 × 10^8^ cells were used following the same procedure. Eluted peptides were analyzed on a Q-Exactive Plus mass spectrometer or an Orbitrap Fusion Tribrid mass spectrometer (Thermo Fisher Scientific, Bremen, Germany). The software Scaffold 4.7.2 (Proteome Software Inc.) was used to validate MS/MS based peptide and protein identifications, perform dataset alignment, and parsimony analysis to discriminate homologous hits. Only proteins identified with more than 95.0% probability (20) and containing at least 2 validated peptides were accepted.

### Cell culture, RNA extraction, RT-PCR, siRNA and plasmid transfections

Human HEK-293 (epithelial), Flp-In T-REx-HEK-293, HeLa (epithelial), DLD-1 (epithelial), Jurkat (T-cell leukemia), U2OS (epithelial osteosarcoma), and MCF-7 (epithelial adenocarcinoma) cells and mouse MEF (fibroblast), C2C12 (myoblast), MEL (erythroleukemia), NS-1 (myeloma), F9 (epithelial carcinoma), Hepa (epithelial hepatoma) cells were grown on plates at 37°C in DMEM with 10% FBS (except DLD-1 and Jurkat cells, which were grown in RPMI).

Total RNA was extracted using the “RNeasy Mini kit” (Qiagen) according to the manufacturer’s recommendations. For gene expression analysis, cDNA synthesis was performed with M-MLV reverse transcriptase (Promega) using 1 μg of total RNA and oligo(dT) primers as described. cDNA for *HCFC2* exons 9–12 was amplified by PCR with sense (GTCAGGATGGACCCTCACAGAC) and antisense (GCCACTGGATTTGAAGGAGTC) primers described in (14).

Complementary siRNA oligonucleotides targeting the *HCFC2* mRNA sequence 5’-GCAAGUCGUUGGUUAUGGA-3’ and Luciferase negative control were purchased from Microsynth. HEK-293 cells were transfected with siRNA duplexes (6 pmol) with Interferin (Polyplus) according to the manufacturer’s instructions. After 24 h an additional transfection was performed. Cells were fixed and used for immunofluorescence 72 h after the initial transfection.

For transient transfection, HEK-293 cells were first seeded onto 10 cm plates in 10 ml of DMEM supplemented with 10% FBS, and 24 h later transfected with 8-10 μg of plasmid combined with 20-30 μl of jetPEI according to the manufacturer’s instructions (Polyplus) and samples collected 24-48 h later.

### Generation of doxycycline-inducible HEK-293 cell lines

To create recombinant HCF-2-encoding Flp-In T-REx-HEK-293 cells, 0.3 μg of pcDNA5FRT/TO vectors containing coding sequences of interest with N-terminal Flag and Cherry tags were co-transfected with 2.7 μg of the Flpase expression vector pOG44 into Flp-In T-REx-HEK-293 cells with jetPEI, and transfected cells were selected in hygromycin-containing medium (100 μg/ml) for 10 days as described (Invitrogen). Synthesis of recombinant proteins was induced with of 1 μg/ml doxycycline in the medium. To inhibit leaky recombinant protein synthesis in the absence of doxycycline, inducible cell lines were maintained in DMEM supplemented with tetracycline-free FBS (BioConcept). Recombinant F-Cherry-HCF-2 proteins were detected by immunoblot with α-Flag and/or α-Cherry antibodies.

### Electrophoretic mobility shift assay (EMSA)

Full-length VP16 (VP16_FL_) and C-terminally truncated VP16 (VP16ΔC) proteins were synthesized using a rabbit TNT quick-coupled transcription/ translation system (Promega) with ^35^S-labeled methionine. The (OCTA+)TAATGARAT ICP0 DNA probe and Oct-1 were prepared as described (21) except that the DNA probe was prepared with fluorescently labeled primers (Microsynth). F-Cherry, F-Cherry-HCF-2_WT_ and F-Cherry-HCF-2_+NLS_ were purified by immunoprecipitation with α-Flag agarose 24 h after induction with doxycycline and WCL extraction from the respective HEK-293 cells. After several washes, recombinant proteins were eluted using Flag-peptide and quantified by immunoblot analysis with α-Cherry antibody. Amounts of VP16 proteins were quantified based on radioactive signal of ^35^S-labeled methionines using a Typhoon TRIO+ imager (Amersham Biosciences) for equal loading. VIC formation and EMSA were as described in (21). The gels were scanned with the Odyssey infrared imager (LI-COR).

### Immunofluorescence

For immunofluorescence, MEF, HEK-293, or HeLa cells were grown at 37°C for 24-48 h on coverslips in 24-well plates, washed with PBS, and fixed with 3.6% formaldehyde for 15 min. Fixed cells were washed three times with PBS containing 0.2% Triton X-100 (PBS-Triton) and further blocked for 30 min at room temperature with 2% goat serum in PBS-Triton, followed by incubation with relevant primary antibodies in the blocking buffer for 1 h. Cells were subsequently washed three times with PBS-Triton and incubated with appropriate secondary antibody (1:400) in the blocking buffer for 30 min. After three washes with PBS-Triton, coverslips were mounted with DAPI-containing Vectashield medium (Reactolab S.A.). Samples were analyzed using a fluorescent (Leica DM 6000 B) or confocal (Zeiss LSM 880) microscope.

### RNA polymerase I (Pol I) inhibition

For Pol I inhibition, cells were treated with actinomycin D (ActD; 0.05 ug/ml or 5 ug/ml), BMH21 (1 uM), or etoposide (25 uM) for 5 h and then immunostained as indicated. Negative control cells were treated with the DMSO vehicle.

### Chromatin and nucleolar purification assays

The small-scale biochemical chromatin fractionation was performed as described (22). Nucleolar purification was performed as described (23). Briefly, 5–10 × 10^6^ HEK-293 or MEF cells were washed with cold sucrose-containing solution I (0.5 M sucrose, 3 mM MgCl_2_ with Cocktail protease inhibitor, Roche), harvested by scraping into 1 ml solution I and sonicated in a BioRuptor for 10 cycles (15 sec ON/15 sec OFF). Disruption of cellular and nuclear membranes was verified by phase contrast microscopy. The suspension was then underlaid with an equal volume of solution II (1 M sucrose, 3 mM MgCl_2_ with Cocktail protease inhibitor) and centrifuged at 1800 g for 5 min at 4°C. The 0.5 M sucrose phase was collected as the cytosolic plus nuclear protein fraction. To purify nucleoli further, the nucleoli-containing pellet was resuspended in 0.5 ml solution I, underlaid with an equal volume of solution II, centrifuged and the pellet resuspended in 1 ml solution I as the nucleolar fraction. Equal amounts of the two fractions were boiled in 1 × Laemmli sample buffer and used for immunoblot analysis.

### Cell counting and metabolic assays

For cell counting, indicated HEK-293 cell lines were seeded onto 6-well plates at 2 × 10^4^ cells per well and 8 h later induced with doxycycline (1ug/ml). Growth curve analysis was started 24 h after doxycycline addition. At the given times, adherent cells were trypsinized and counted using a NucleoCounter NC-250 machine (Chemometec). For metabolic assays, indicated HEK-293 cells (1000 cells/well) were seeded onto 96-well plates (one plate per time point) in replicates of five per assay. Recombinant protein synthesis was induced by adding doxycycline (1ug/ml) after 8 h and metabolic rates measured by the MTT assay (reducing of tetrazolium dye 3-(4,5-dimethylthiazol-2-yl)-2,5-diphenyltetrazolium bromide) over the next eight days as recommended by the manufacturer.

### RNA-seq

Poly (A)-containing mRNA from each sample was used to prepare libraries using NEXTflex qRNA-seq Kit for 125 nucleotide single-end sequencing by Illumina HiSeq 2500. Reads were mapped onto the human genome (GRCh37, version 19 (Ensembl 74) and its corresponding GENCODE annotation file, version 2013-12-05) using STAR with the outFilterMatchNminOverLread parameter set to 0.4. Expression values are given as fragments per kilo base per million mapped reads (FPKM) and the expected counts per transcript were then calculated using rsem-calculate-expression from RSEM-1.3.0 software. Only mRNAs with an FPKM above 1 were kept for further analysis.

### Bioinformatic analyses

The initial Minimum Evolution (ME) tree for the HCF-protein Kelch-domain was built in MEGA X (24) using Neighbor joining (25). Distances were computed using the Dayhoff matrix based method (26). The ME tree was searched using the Close-Neighbor-Interchange algorithm (27).

Differential gene expression analysis was performed with DESeq2 (28). A comparison of gene expression levels was done separately for each HCF-2_WT_ and HCF-2_Fn3nc*_ sample with respect to day 1. Genes were considered differentially expressed if the log2 of fold expression change was greater than 0.5 or less than −0.5 with a False Discovery Rate (FDR) of < 0.05. This analysis led to the identification of 7175 differentially expressed genes.

For differential gene expression clustering, we used the partitioning around medoid (PAM) algorithm from the R‘cluster 1.14.4’ library on the 7175 gene set using 1 minus the Pearson correlation coefficient as the dissimilarity measure. The DESeq2 normalized counts were centered and scaled to obtain z-scores. The clustering results were displayed using the ‘ggplot 2.14.1’ R library heatmap.2 function.

Gene Ontology (GO) terms enrichment analysis was done with the clusterProfiler R package (29) and corroborated with goseq (30). The REVIGO on-line service was used to reduce semantic redundancy of lists of enriched GO terms produced by clusterProfiler (31). Gene Set Enrichment Analysis (GSEA) was performed as described in (32).

## RESULTS

### HCF-2 arose with vertebrate evolution

Aiming to gain insights into functional divergence between HCF-1 and HCF-2, we first compared the previously reported domain organization of the two proteins (14, 17, 33). Figure 1A shows a comparison of human HCF-1 with its human and mouse HCF-2 orthologs. HCF-1 consists, from N-to C-terminus, of a Kelch domain, which tethers HCF-1 to chromatin; a half fibronectin type 3 repeat (Fn3n); regions enriched in basic (Basic) or acidic (Acidic) amino acids, separated by the HCF-1_PRO_ repeats; one-and-a-half Fn3 repeats (Fn3c); and a nuclear localization signal (NLS). After OGT proteolytic processing, the resulting N- and C-terminal fragments remain non-covalently associated via a two Fn3-repeat module, called here Fn3nc, created by the half Fn3n repeat and one-and-half Fn3c repeats (21). Together this interdigitated Fn3 self-association structure has been referred to as SAS1 for “self-association sequence 1” (34); for clarity, here we refer to this SAS1 element as the Fn3nc module. Although many HCF-1 elements, such as Basic, HCF-1_PRO_ repeat, Acidic and NLS elements are missing in HCF-2, the remaining human Kelch (68% identical), Fn3n (51% identical), and Fn3c (56% identical) modules are highly related between these two paralogs (14).

**Figure 1.**
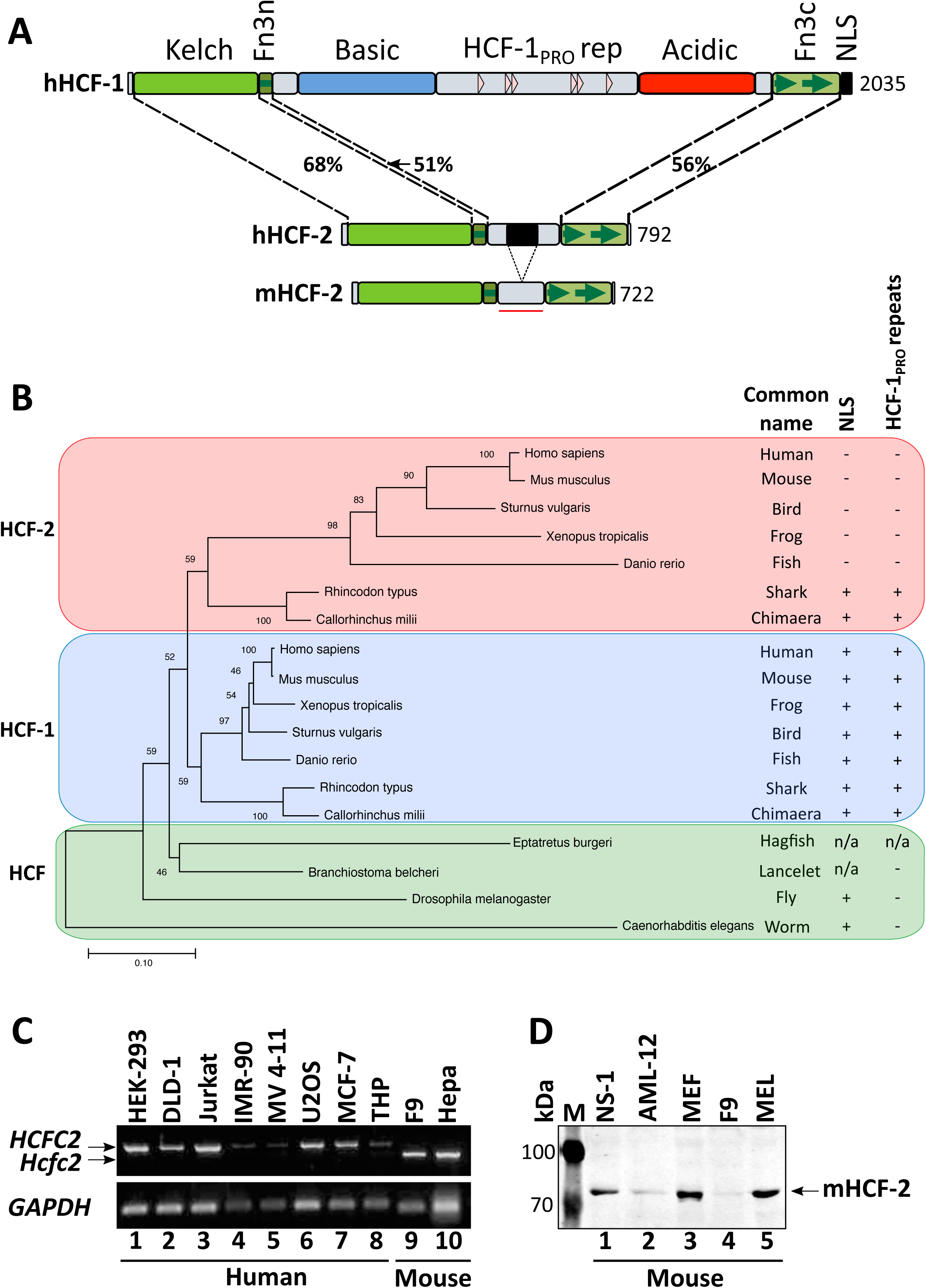
HCF-2 conservation in is vertebrates and cognate gene expression in mammalian cells. ***(A)*** Diagrams of human HCF-1, and human and mouse HCF-2. Percentage identity between the human HCF-1 and HCF-2 Kelch, Fn3n, and Fn3c segments is given. NLS, Nuclear localization signal. The Fn3n-to-Fn3c linker region in mHCF-2 used for antibody generation is indicated with a red bar. ***(B)*** Evolutionary tree of HCF-protein Kelch domains from 11 animal species inferred using the Minimum Evolution method (58). The optimal tree with the sum of branch length = 2.66193529 is shown. The percentage of replicate trees in which the associated taxa clustered together in the bootstrap test (500 replicates) (59) are shown next to the branches. The tree is drawn to scale, with branch lengths in the same units as those of the evolutionary distances used to infer the phylogenetic tree. Common name of species, presence of NLS and HCF-1_PRO_ repeats are shown separately. n/a, not available. ***(C)*** RT-PCR of *HCFC2* in the indicated cell lines of human (lanes 1–8) and mouse (lanes 9–10) origin. ***(D)*** Immunoblot of the indicated mouse cell line extracts probed with α-HCF-2 antibody described in Supplementary Figure S1.

To study the evolution of HCF-encoding genes, we focused on the Kelch domain for two reasons: (i) it is the most highly conserved region in HCF proteins and (ii) we did not always find the C-terminal Fn3c coding sequences in the genome of early chordates (e.g., hagfish). Figure 1B shows an evolutionary tree of Kelch domain sequences (amino acids 17–359 in human HCF-1) from two invertebrate non-chordate species (worm and fly), one non-vertebrate chordate (lancelet), and one jawless (hagfish) and seven jawed– two cartilaginous (chimaera and shark) and five bony (fish, amphibian, bird, mouse and human) – vertebrates. Only one HCF-like protein was observed in the four non-jawed species, whereas all jawed vertebrate species had two HCF-like proteins, which segregated into HCF-1- and HCF-2 Kelch-domain-related groups. Also segregating with the HCF-1, but not the HCF-2, Kelch domain in bony vertebrates are the Basic and Acidic regions, the central HCF-1_PRO_ proteolytic processing repeats and C-terminal NLS (Figure 1B). The cartilaginous vertebrate HCF-2 proteins are an exception as discussed further below.

In mouse and human, HCF-2 proteins are encoded by intron-containing genes on different chromosomes from the HCF-1 protein-encoding genes; the corresponding *Hcfc1*/*HCFC1* genes are X-linked and *Hcfc2*/*HCFC2* genes are autosomal (Chr10 in mouse and Chr12 in human). These observations suggest that HCF-2 arose via a gene duplication at the time of jawed vertebrate evolution.

### HCF-2 is broadly present in human and mouse cells

HCF-2-encoding mRNA is broadly present in human tissues (14). Here, as shown in Figure 1C, we probed HCF-2-encoding mRNA levels in tissue-culture cells by RT-PCR. We detected HCF-2-encoding transcripts in all eight human and two mouse cell lines tested. DNA sequence analysis showed that the mouse PCR product lacks the sequence corresponding to human exon 11, which encodes a 67 aa segment lying between the Fn3n and Fn3c elements as shown in Figure 1A (bottom). This segment is missing in both rat and mouse HCF-2 (Supplementary Figure S1A) and the corresponding *HCFC2* genes contain mutations that lead to exon deletion (rat) or likely exon skipping (mouse). These results suggest that the non-conserved sequences between the HCF-2 Fn3n and Fn3c segments are not functionally critical.

To study the native HCF-2 protein, we generated an affinity purified anti-mouse HCF-2 (α-HCF-2) rabbit polyclonal antiserum (see Materials and Methods, and Supplementary Figure S1 for details). We probed cell extracts from five mouse cell lines by immunoblot and detected HCF-2 protein in each case, indicating its broad presence in cell lines (Figure 1D; see also Supplementary Figure S1E). Although this anti-mouse HCF-2 antibody was less effective in detecting human HCF-2 in immunoblots (Supplementary Figure S1F), it detected human HCF-2 via immunoprecipitation (Supplementary Figure S1B) and immunofluorescence (see below). These results, combined with the broad presence of HCF-2-encoding mRNAs, suggest that HCF-2 is a common protein that can participate in the general functions of mammalian cells.

### HCF-2 fails to form a VIC with full-length VP16

HCF-1 forms the VIC with VP16 through the activity of the Kelch domain (14, 35–37), and C-terminal HCF-1 NLS (21, 35). VIC formation, however, with a truncated VP16 protein, which lacks its C-terminal acidic transcriptional activation domain (called VP16ΔC) does not require the HCF-1 NLS, and both HCF-1 and HCF-2 can form a VIC with VP16ΔC via the conserved Kelch domain. As bony vertebrate HCF-2 proteins lack an evident NLS (Figure 1B), we asked whether human HCF-2 can form a complex with VP16_FL_ in an EMSA. As expected, in the absence of any HCF protein neither VP16_FL_ nor VP16ΔC formed a VIC, even though the Oct-1 DNA-binding POU domain bound to the VIC-forming DNA probe (Figure 2A, lanes 1-3). In contrast, the epitope-tagged F-Cherry-HCF-2_WT_ purified from HEK-293 cells, readily formed a VIC with VP16ΔC, but not with native VP16_FL_ (Figure 2A, lanes 4-6), indicating that HCF-2 is not likely a target of VP16 during HSV infection. Interestingly, the mere addition of the HCF-1 NLS to the C-terminus of HCF-2 was sufficient to promote VIC formation with VP16_FL_ (Figure 2B, lanes 1-3). These results suggest that HCF-2 owing to the absence of an NLS, does not directly influence HSV infection, but has retained the ability to interact with some targets of the HCF-1 Kelch domain (illustrated here with VP16ΔC).

**Figure 2.**
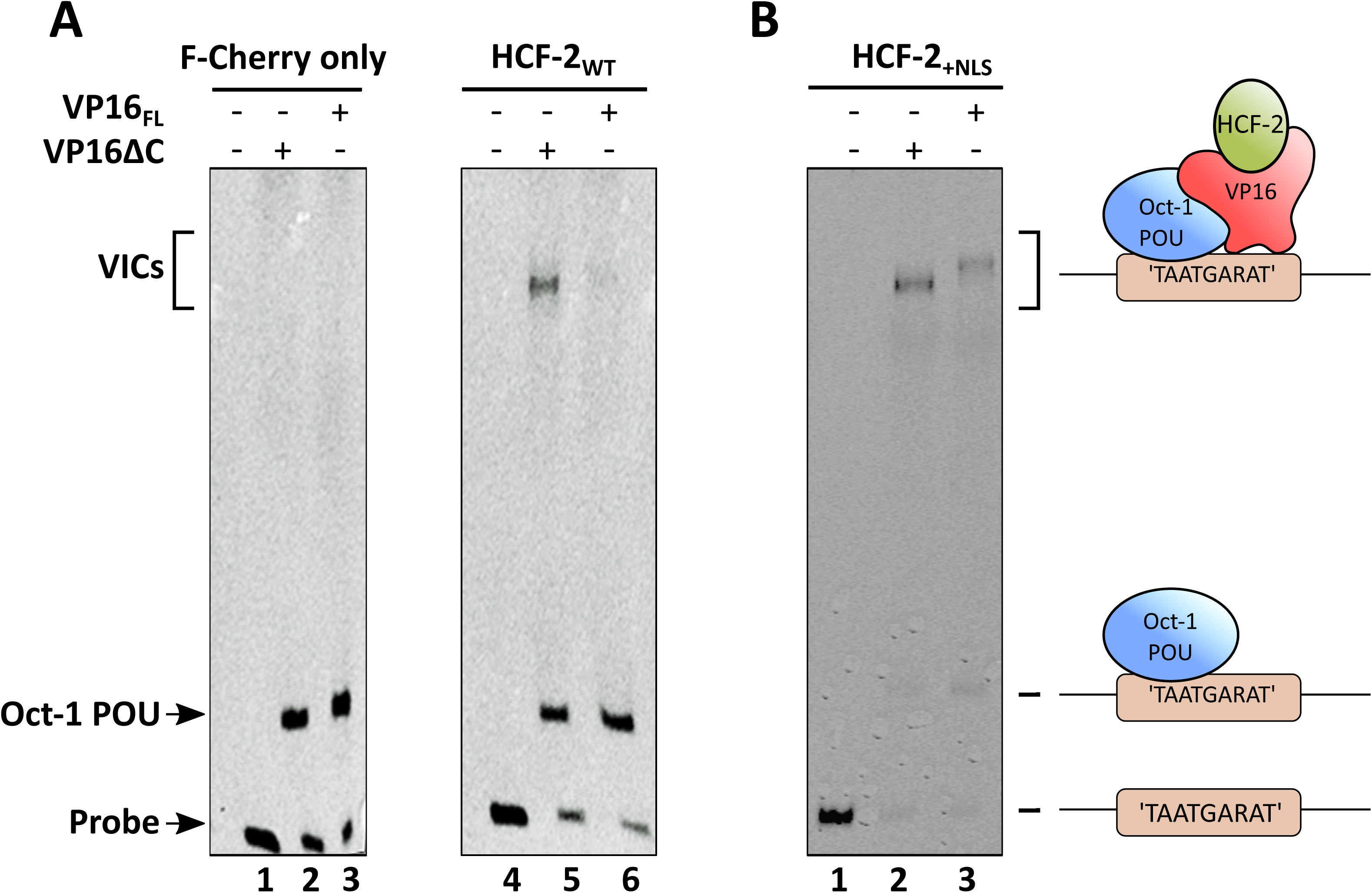
HCF-2 fails to form a VIC with VP16_FL_ ***(A)***, but addition of HCF-1 NLS sequence to the C-terminus of HCF-2 activates VIC formation with VP16_FL_ ***(B)***. Immunopurified F-Cherry, F-Cherry-HCF-2_WT_ or F-Cherry-HCF-2_+NLS_ proteins were used for EMSA assay (see Materials and Methods). Lanes 1 and 4, probe alone with Oct-1 POU domain; lanes 2 and 5, Oct-1 POU domain and VP16ΔC; lanes 3 and 6, Oct-1 POU domain and VP16_FL_. ***(A)*** and (***B)*** represent two different gels; models of VIC composition and formation are shown.

### HCF-1 and HCF-2 display different subnuclear localizations

In principal, because the Fn3n element of HCF-2 has been shown to bind the Fn3c element of HCF-1 (34), HCF-1 and HCF-2 can associate with each other in the cell via the conserved HCF Fn3n and Fn3c interaction. Having antibodies against both endogenous HCF-1 and HCF-2, we therefore asked whether immunoprecipitation with either one or the other HCF-1 or HCF-2 antibody leads to recovery of both HCF-1 and HCF-2 thus indicating their association. Figure 3A indicates that HCF-1 and HCF-2 do not obviously associate with one another as, after separate HCF-1 and HCF-2 immunoprecipitation from a MEF-cell lysate, only HCF-1 is evident from the HCF-1 immunoprecipitate (Figure 3A, upper panel, compare lanes 1 and 2) and only HCF-2 is evident from the HCF-2 immunoprecipitate (lower panel).

**Figure 3.**
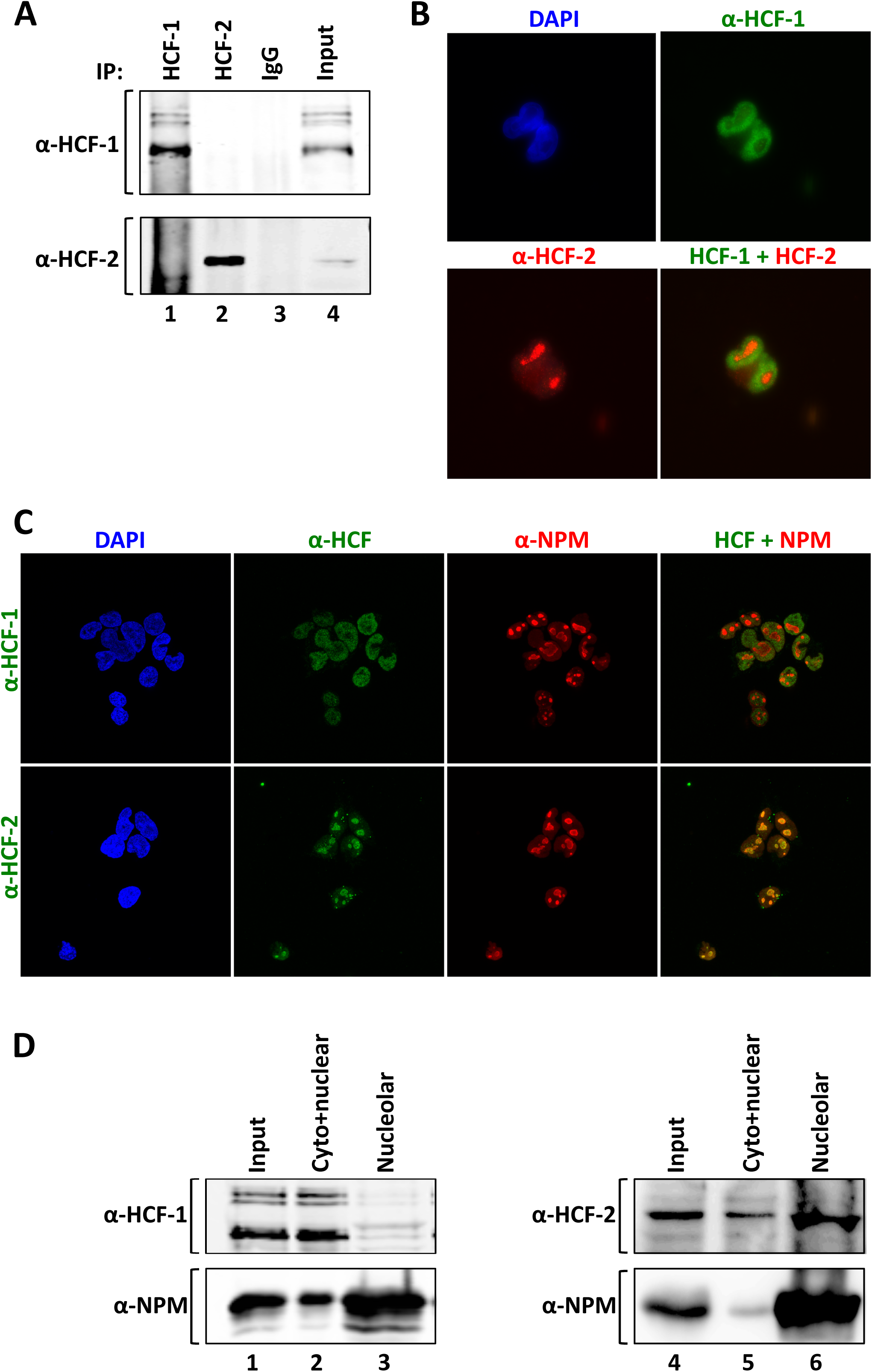
HCF-1 and HCF-2 display different subnuclear localization. ***(A)*** α-HCF-1 (top) and α-HCF-2 (bottom) immunoblot of protein complexes after immunoprecipitation from a MEF whole-cell extract with α-HCF-1 antibody (lane 1), α-HCF-2 antibody (lane 2), or non-immune IgG (lane 3). Lane 4, starting whole-cell lysate. ***(B)*** HEK-293 cells co-stained with mouse monoclonal α-HCF-1 (green) and rabbit polyclonal α-HCF-2 (red) antibodies. Left, nuclear staining with DAPI (blue); right, merge of individual HCF-1 and HCF-2 signals. ***(C)*** Co-immunostaining of HEK-293 cells with α-NPM antibody (red) and either polyclonal rabbit α-HCF-1 antibody (green, upper panels) or α-HCF-2 antibody (green, lower panels). Left panels, nuclear staining with DAPI (blue). Right, merge of NPM and HCF signals. ***(D)*** Immunoblot of samples after nucleolar fractionation of HEK-293 cells with HCF-1, HCF-2, and NPM antibodies as indicated. Lanes 1 and 4, starting whole-cell lysate; lanes 2 and 5, cytoplasmic and non-nucleolar fraction; lanes 3 and 6, nucleolar fraction.

This result led us to compare directly the cellular localization of endogenous HCF-1 and HCF-2. Immunostaining of endogenous HCF-1 in cultured cells generates a clear nuclear pattern (33, 34), owing in part to its NLS (22, 38). Although mammalian HCF-2 proteins lack an evident NLS, immunostaining of endogenous HCF-2 in mouse fibroblasts revealed nuclear staining but with a punctate pattern (Supplementary Figure S2A) — a staining pattern that was reduced when cells were pretreated with siRNA against HCF-2-encoding mRNA indicating an HCF-2-specific signal (Supplementary Figure S2B). This HCF-2 pattern was clearly distinct from that generated for HCF-1 as shown by the non-overlapping co-immunostaining of human HEK-293 cells with a mouse monoclonal α-HCF-1 antibody (18) and the rabbit polyclonal HCF-2 antibody (Figure 3B). Thus, HCF-1 and HCF-2 do not appear to associate in cultured cells under normal conditions and possess different nuclear localization properties.

The punctate nuclear HCF-2 staining pattern was reminiscent of nucleoli, the site of ribosome synthesis, with the HCF-1 pattern representing a complementary non-nucleolar pattern. To analyze these different nuclear localization properties further, we compared the HCF-1 and HCF-2 staining patterns with the nucleolar marker nucleophosmin (NPM, also known as B23) in HEK-293 cells. Indeed, consistent with the aforementioned nucleolar vs. non-nucleolar HCF-2/HCF-1 staining patterns, as shown in Figure 3C, the HCF-2 pattern co-localized with the nucleolar NPM marker (lower panel), whereas HCF-1 did not (upper panel).

Consistent with HCF-2 nucleolar localization, the ribosomal RNA (rRNA) synthesis inhibitors ActD and BMH21, which induce nucleolar disassembly, but not the apoptosis inducer etoposide, disrupted the punctate HCF-2 staining (Supplementary Figure S2C).

The different localization of HCF-1 and HCF-2 was substantiated by biochemical fraction. In previous biochemical studies of subnuclear HCF-1 localization, we have used a small-scale chromatin isolation method (39) to probe HCF-1 chromatin-binding properties. This chromatin fractionation scheme, however, as shown in Supplementary Figure S3A and S3B, does not separate nucleolar components (e.g., NPM, NCL, and RPA194) from the chromatin (e.g., HCF-1 and histone H3) fraction. We therefore turned to a small-scale nucleolar isolation method (23). Immunoblot analysis of cytosol- (MEK and tubulin), chromatin- (histone H3), soluble nucleoplasm- (sc-35) and nucleolus- (NPM) specific markers demonstrates the successful fractionation of nucleoli (Supplementary Figure S3C). Using this fractionation scheme, HCF-1 fractionates primarily with the non-nucleolar ‘cyto+nuclear’ fraction (Figure 3D, compare lanes 2 and 3), whereas HCF-2 fractionates primarily in the nucleolar fraction with NPM (lanes 5 and 6), further emphasizing the different subnuclear localization of these two evolutionarily related proteins.

### HCF-2 is localized in the nucleolar fibrillar center

The nucleolus is primarily involved in ribosome biogenesis and is organized into three compartments (Figure 4A): (i) the fibrillar center (FC), containing RNA polymerase I (Pol I) and its co-regulators; (ii) the dense fibrillar component (DFC) representing proteins and small nucleolar RNA’s (snoRNA) involved in rRNA maturation; and (iii) the granular component (GC) where pre-ribosomal particles are assembled (40, 41). These three compartments reflect the tripartite role of the nucleolus in ribosome synthesis: rRNA synthesis, maturation, and subsequent ribosome assembly (42).

**Figure 4.**
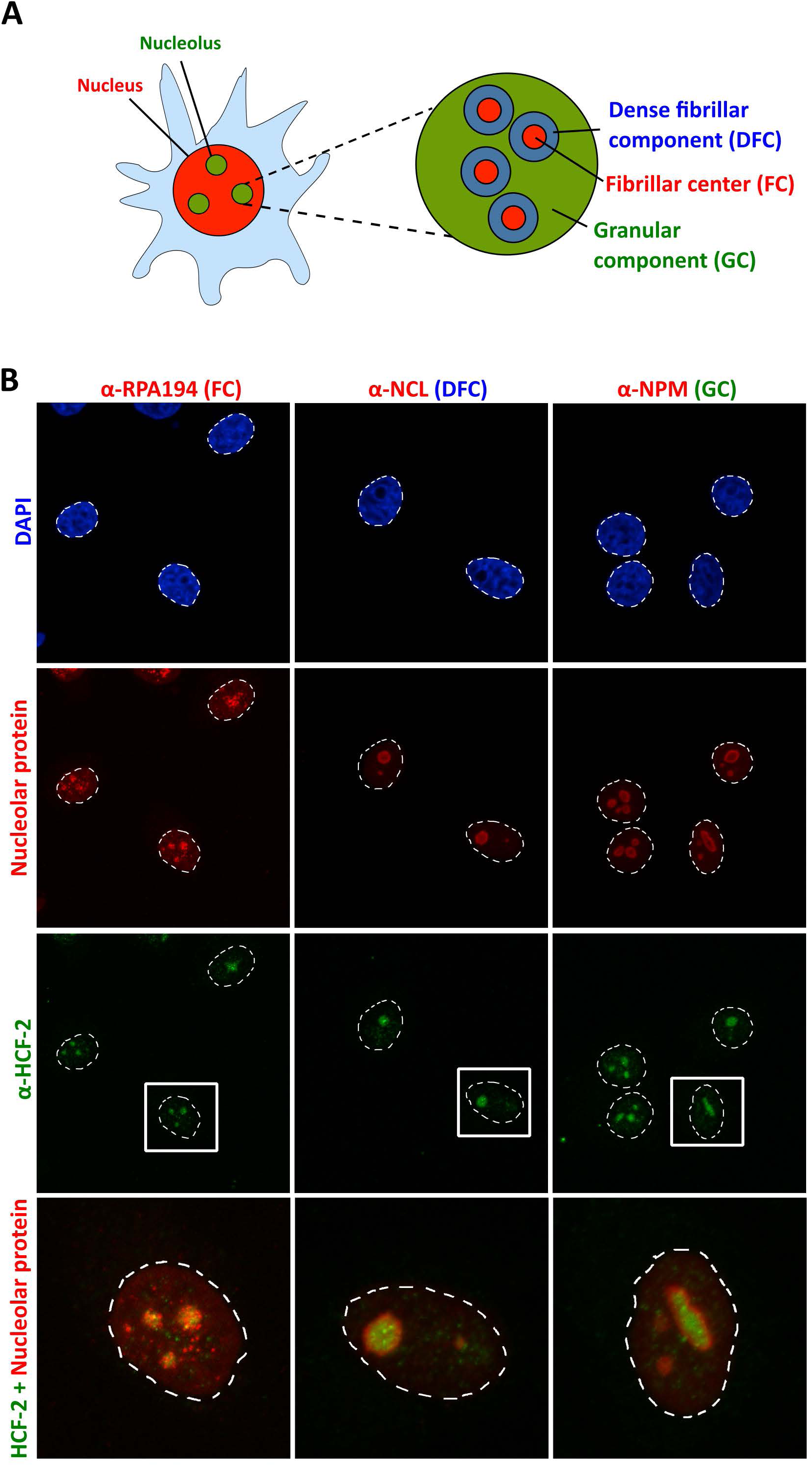
HCF-2 is localized in the fibrillar center of nucleolar compartment. ***(A)*** Diagram of mammalian nucleolus with fibrillar centers (FC, red) surrounded by dense fibrillar component (DFC, blue) and further encircled by the granular component (GC, green). ***(B)*** Co-immunostaining of HeLa cells with α-HCF-2 antibody (green) and (left to right) antibodies for markers for the different subnucleolar regions (red): RPA194 (FC), NCL (DFC), and NPM (GC). Upper panels, nuclear staining with DAPI (blue). Bottom panels, enlargement of one cell with merge of HCF-2 and nucleolar maker signals is shown. Dashed line denotes the nuclear area.

To determine in which subnucleolar region HCF-2 resides, we performed co-immunostaining of HCF-2 and nucleolar proteins corresponding to the FC with the Pol I subunit RPA194, the DFC with nucleolin (NCL); and the GC with NPM (Figure 4B). The HCF-2 signal clearly overlapped with the FC Pol I RPA194 signal.

Thus, despite lacking the canonical NLS present in HCF-1, HCF-2 is still imported into the nucleus and more specifically into nucleoli.

### HCF-2 association with NPM

The importation of proteins into the nucleolus is often promoted by one of two nucleolar shuttle proteins – NPM and NCL (43). In an MS analysis of proteins recovered in an endogenous HCF-2 immunoprecipitate from MEF cells, we identified NPM, but not NCL, as a directly or indirectly associating HCF-2 protein (see Supplementary Table S1). An HCF-2–NPM association was further supported by immunoblot analysis of a separate α-HCF-2 immunoprecipitation (Supplementary Figure S4). Perhaps NPM plays a role in HCF-2 nucleolar importation but if so after importation NPM would probably dissociate from HCF-2 as NPM and HCF-2 do not obviously co-localize in the nucleolus (see Figure 4B).

### The Fn3c element promotes HCF-2 nucleolar localization

Unlike a nucleus, which is enclosed in a double membrane and where the presence of a clearly defined NLS in a protein can promote its nuclear entry, the nucleolus has no surrounding membrane and, although several amino acid sequences have been considered as nucleolar localization sequences (NoLS) (43–46), there is no well-defined NoLS (43, 47, 48).

To identify the region of HCF-2 responsible for nucleolar targeting, we prepared an N-terminal Flag and fluorescent Cherry tag–HCF-2 fusion (called F-Cherry–HCF-2) under the control of a doxycycline-inducible promoter (Figure 5A, compare lanes 4 and 7) and made a panel of stable HEK-293 cell lines with various ectopic truncated HCF-2 proteins. Biochemical nucleolar fractionation showed that, in contrast to the F-Cherry tag alone, ectopically synthesized tagged full-length HCF-2 is present, albeit not entirely, in the nucleolar-enriched fraction (compare lanes 7–9 with 1–3); immunofluorescence of these cells showed that the non-nucleolar F-Cherry-HCF-2 is primarily nuclear (Supplementary Figure S5A). An initial truncation analysis showed that the combined N-terminal HCF-2 Kelch-domain and Fn3n sequences were neither important nor sufficient for nucleolar localization (data not shown) — indeed, the HCF-2 Fn3c element alone promotes F-Cherry-HCF-2 nucleolar localization (lanes 13–15).

**Figure 5.**
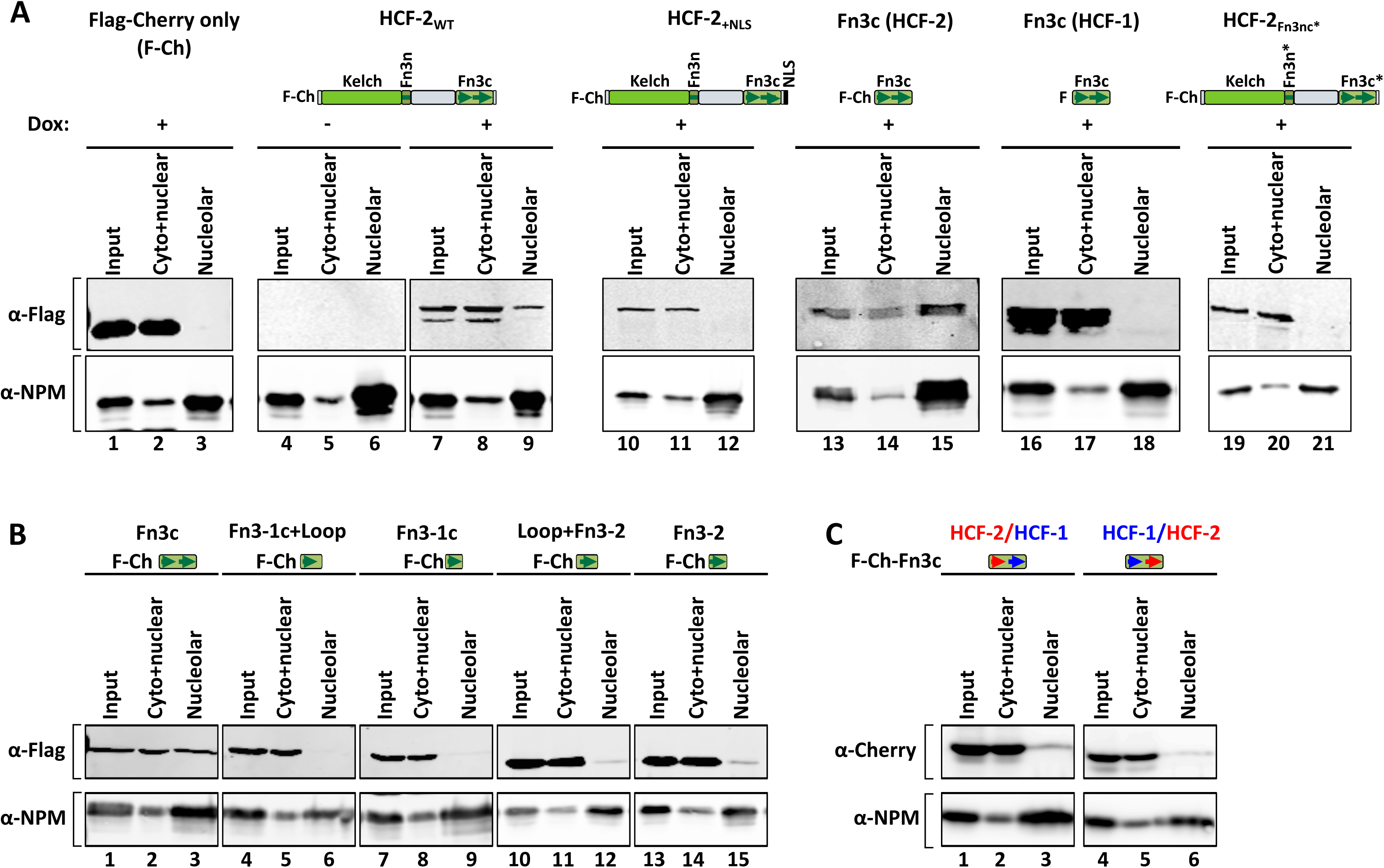
Fn3c facilitates HCF-2 but not HCF-1 nucleolar localization. **(*A*)** F-Cherry-only (F-Ch), F-Cherry-HCF-2_WT,_ F-Cherry-HCF-2_+NLS_, F-Cherry-HCF-2_Fn3c_, F-HCF-1_Fn3c_ or F-Cherry-HCF-2_Fn3nc*_ (see respective protein structure models on top) synthesis in HEK-293 T-REx cells was induced with doxycycline (1 uM for 24 h) and samples were subjected to nucleolar fractionation. Induction of recombinant protein synthesis is shown for F-Cherry-HCF-2_WT_ (–, without doxycycline; +, with doxycycline). Presence or absence of recombinant HCF proteins was analyzed with α-Flag antibody (upper panels), and the quality of nucleolar purification assayed with the α-NPM antibody (lower panels). **(*B*)** Both Fn3c type repeats facilitate nucleolar localization of HCF-2. HEK-293 cells were transiently transfected with plasmids encoding F-Cherry-HCF-2_Fn3c_ truncations and 24 h later collected for nucleolar fractionation. Presence or absence of recombinant F-Cherry-HCF-2_Fn3nc_ truncations was analyzed with α-Flag antibody (upper panels), and the quality of nucleolar purification assayed with the α-NPM antibody (lower panels). **(*C*)** As in **(*B*)**, but with plasmids encoding hybrid HCF-1 and HCF-2 F-Cherry-Fn3c sequences. HCF-2/HCF-1, Fn3-1c_HCF-2_/Fn3-2_HCF-1_ hybrid; HCF-1/HCF-2, Fn3-1c_HCF-1_/Fn3-2_HCF-2_.

As aforementioned, the Fn3nc module consists of two Fn3 repeats: a hybrid Fn3-1n and Fn3-1c repeat, and Fn3-2; the two Fn3 repeats are linked by a highly conserved 6 aa loop (see Supplementary Figure S5B). With SwissModel visualized by PyMOL, we used the HCF-1 Fn3nc structure (21) to build a model of HCF-2 Fn3nc; Supplementary Figure S5C shows comparisons of the determined HCF-1 and deduced HCF-2 structures. Despite over 40% differences in the HCF-1 and HCF-2 Fn3nc amino acid sequences (Figure 1A, Supplementary Figure S5B), HCF-2 has the potential to form a 3D Fn3nc structure similar to that of HCF-1, reinforcing the conclusion that the Fn3nc element has been conserved in the HCF-protein family.

To further define the HCF-2 Fn3c NoLS, we created a set of HCF-2 Fn3c truncation mutants which carried only the Fn3-1c or Fn3-2 repeat with or without the loop (Figure 5B). Compared to the entire Fn3c element (Figure 5B, lanes 1–3), all of the Fn3c truncations were less abundant in the nucleolar fraction, with Fn3-2 truncations essentially absent (lanes 4–9) and Fn3-1c truncations reduced (lanes 10–15). Thus, HCF-2 nucleolar localization appears to depend on either multiple sequence elements or one sequence element that incorporates parts of both Fn3-1c and Fn3-2.

Although Fn3c is sufficient for nucleolar localization, in full-length HCF-2 nucleolar localization may also depend on Fn3n and Fn3c association. Park et al (21) have shown that the W384A Fn3n substitution and V1866E Fn3c substitution in HCF-1 — two residues buried within the Fn3n-Fn3c interface of the Fn3nc structure — disrupt formation of the Fn3nc element. We therefore introduced the respective mutations (W377A and V653E) into HCF-2 (creating HCF-2_Fn3nc*_) to test whether disruption of the Fn3nc module might affect HCF-2 nucleolar localization. As Figure 5A (lanes 19-21) shows, these mutations indeed disrupt nucleolar localization. Thus, curiously, removal of the HCF-2 Kelch and Fn3n regions does not prevent HCF-2 Fn3c nucleolar localization, but their retention, in a mutant form that likely prevents Fn3nc module formation, prevents nucleolar localization. Alternatively, the V653E Fn3c mutation itself directly disrupts nucleolar localization.

### The HCF-1 Fn3c element lacks nucleolar localization properties

Consistent with the lack of HCF-1 nucleolar localization (see Figure 3B and 3C), unlike the HCF-2 Fn3c element, a Flag-tagged HCF-1 Fn3c element is not nucleolar (Figure 5A, lanes 16–18), suggesting that differing HCF-1 and HCF-2 subnuclear distribution is in part defined by differences in their respective Fn3c elements. To investigate these differences further, we prepared two chimeric HCF-1 and HCF-2 mutants by swapping Fn3-1c and Fn3-2 repeats (Figure 5C). Although diminished, both chimeric proteins were still evident in the nucleolar fraction (compare the nucleolar recovery of HCF-2 and HCF-1 Fn3c in Figure 5A, lanes 13–18, with that of the two chimeras in Figure 5C). These results are consistent with the involvement of both HCF-2 Fn3-1c and Fn3-2 repeats in nucleolar localization. We thus suggest that HCF-2 nucleolar localization is an acquired activity that arose after HCF-gene duplication — an acquisition involving changes in both the Fn3-1c and Fn3-2 repeats.

### The HCF-1 NLS inhibits HCF-2 nucleolar localization

In Figure 2B, we showed that the HCF-1 NLS can stimulate VIC formation when fused to HCF-2. Here, we asked whether the HCF-1 NLS would affect HCF-2 cellular localization. Indeed, the aforementioned HCF-2_+NLS_ molecule — capable of VIC formation — no longer segregates in the nucleolar fraction upon biochemical fractionation (Figure 5A, lanes 10-12). These results suggest that loss of an NLS was an important part of the evolution of the HCF-2 proteins vis-à-vis both its viral-host and cellular activities.

### Ectopic HCF-2 inhibits cell growth and promotes mitotic defects

HCF-1 is known to regulate cell growth and division — cells lacking HCF-1 become G1-phase arrested and develop multiple mitotic defects (e.g., micronuclei and multi-nucleation) (11–13). Having stable inducible HEK-293 F-Cherry-HCF-2 cell lines, we tested the effect of elevated HCF-2 levels on cell growth and division after HCF-2 induction with doxycycline. Strikingly, cells with elevated levels of full-length F-Cherry-HCF-2 (nucleolar and nuclear, Figure 5A) have slower growth rates (as assayed by cellular mitochondrial activity in Supplementary Figure S6A and by cell counting in Figure 6A) compared to cells with elevated F-Cherry protein levels. Notably, the HCF-2_Fn3nc*_ and HCF-2_NLS_ mutants were less capable of inhibiting proliferation (Figure 6A), suggesting that HCF-2 inhibition of cell proliferation is a bona fide property of HCF-2 in these cells.

**Figure 6.**
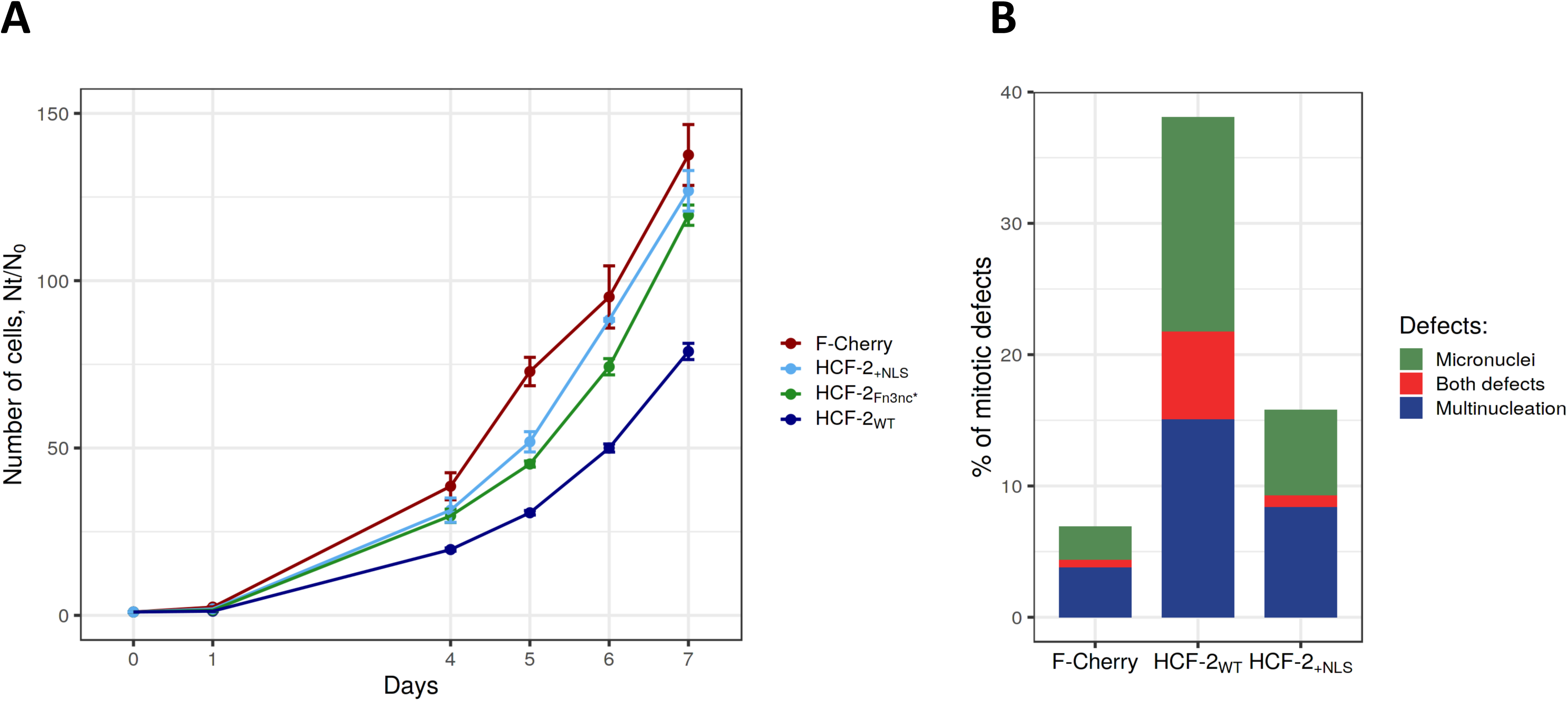
Ectopic HCF-2 synthesis affects cell proliferation **(*A*)** and promotes mitotic defects **(*B*)**. **(*A*)** Growth curves of HEK-293 T-REx cells with induced F-Cherry, F-Cherry-HCF-2_WT,_ F-Cherry-HCF-2_+NLS_, or F-Cherry-HCF-2_Fn3nc*_ synthesis during 7 days after doxycycline addition at day 0. **(*B*)** Percentage of cells with mitotic defects (micronuclei, green; multinucleation, blue; or both, red) in F-Cherry, F-Cherry-HCF-2_WT_ and F-Cherry-HCF-2_+NLS_ containing cells 72 h after doxycycline induction.

Analysis of multinucleation and micronuclear formation (Supplementary Figure S6B) of F-Cherry-HCF-2 cells at 72 h post doxycycline induction revealed a 4-5-fold increase in cell multinucleation (to about 22% of cells) compared to 4% in the F-Cherry-only control (Figure 6B). In contrast, F-Cherry-HCF-2_+NLS_ cells displayed a reduced frequency of these two mitotic defects as compared to wild-type F-Cherry-HCF-2 (Figure 6B).

Together, these results indicate that elevated ectopic HCF-2 levels lead to phenotypes observed when HCF-1 function is lost: reduced proliferation, and mitotic and cytokinesis defects.

### Elevated HCF-2 levels activate differentiation and morphogenesis gene expression programs

Given that elevated HCF-2_WT_, but not HCF-2_Fn3nc*_, levels inhibit cell proliferation, we compared the effects of elevated HCF-2_WT_ and HCF-2_Fn3nc*_ levels on gene expression, by performing a high-throughput RNA-sequence (RNA-seq) time-course analysis of poly(A)-selected RNAs from HCF-2_WT_ and HCF-2_Fn3nc*_ HEK 293 cells (Supplementary Table S2). Ectopic HCF-2_WT_ or HCF-2_Fn3nc*_ synthesis was induced in duplicate with doxycycline and samples collected 1, 2, 4, and 6 days post induction (16 samples total; Supplementary Figure S7).

In the RNA-seq analysis, we could distinguish levels of endogenous from ectopic HCF-2-encoding mRNAs because only the mRNAs of endogenous origin carry native 3’ untranslated region (3’UTR) sequences. As shown in Supplementary Figure S7A, the levels of induced ectopic HCF-2_WT_ or HCF-2_Fn3nc*_ mRNAs were already very elevated by day 1.

A combined principal component analysis (PCA) of the 16 individual HCF-2_WT_ and HCF-2_Fn3nc*_ RNA-seq results revealed in both cases a linear progression over time with duplicate samples positioned in very close proximity to each other, indicating that the time course and RNA-seq analysis were robust (Supplementary Figure S7B). Interestingly, however, the initial day 1 positions and the directions of the sample progressions differed. The difference in initial day 1 positions is likely owing to leaky expression of the HCF-2_WT_ and HCF-2_Fn3nc*_ vectors in uninduced cells (data not shown); the different directions of sample progression probably reflect the induction of different gene-expression programs.

Consistent with the PCA, the maximal gene-expression changes over time were on day 6 in each case, with more genes up-regulated (2199 vs. 1241) in the HCF-2_WT_ samples and more genes down-regulated (2513 vs. 1555) in the HCF-2_Fn3nc*_ samples after six days compared to day 1 (Supplementary Figure S8A). Notably, there was little overlap between the genes affected by the induced synthesis of HCF-2_WT_ or HCF-2_Fn3nc*_ (Supplementary Figure S8B), indicating that the HCF-2_Fn3nc*_ mutation seriously disrupts HCF-2 function.

In total, we identified 7175 genes that were differentially expressed after day 1 (see Materials and Methods for cutoff) in any of the HCF-2_WT_ or HCF-2_Fn3nc*_ samples. The results of partitioning around medoids (PAM) clustering of the expression patterns of these 7175 genes into four clusters (called I-IV) followed by heat-map visualization are shown in Figure 7A and 7B. Figure 7C shows the fold change in transcript abundance relative to day 1 for each gene (ordered as in Figure 7A) at each timepoint for both the HCF-2_WT_ and HCF-2_Fn3nc*_ samples.

**Figure 7.**
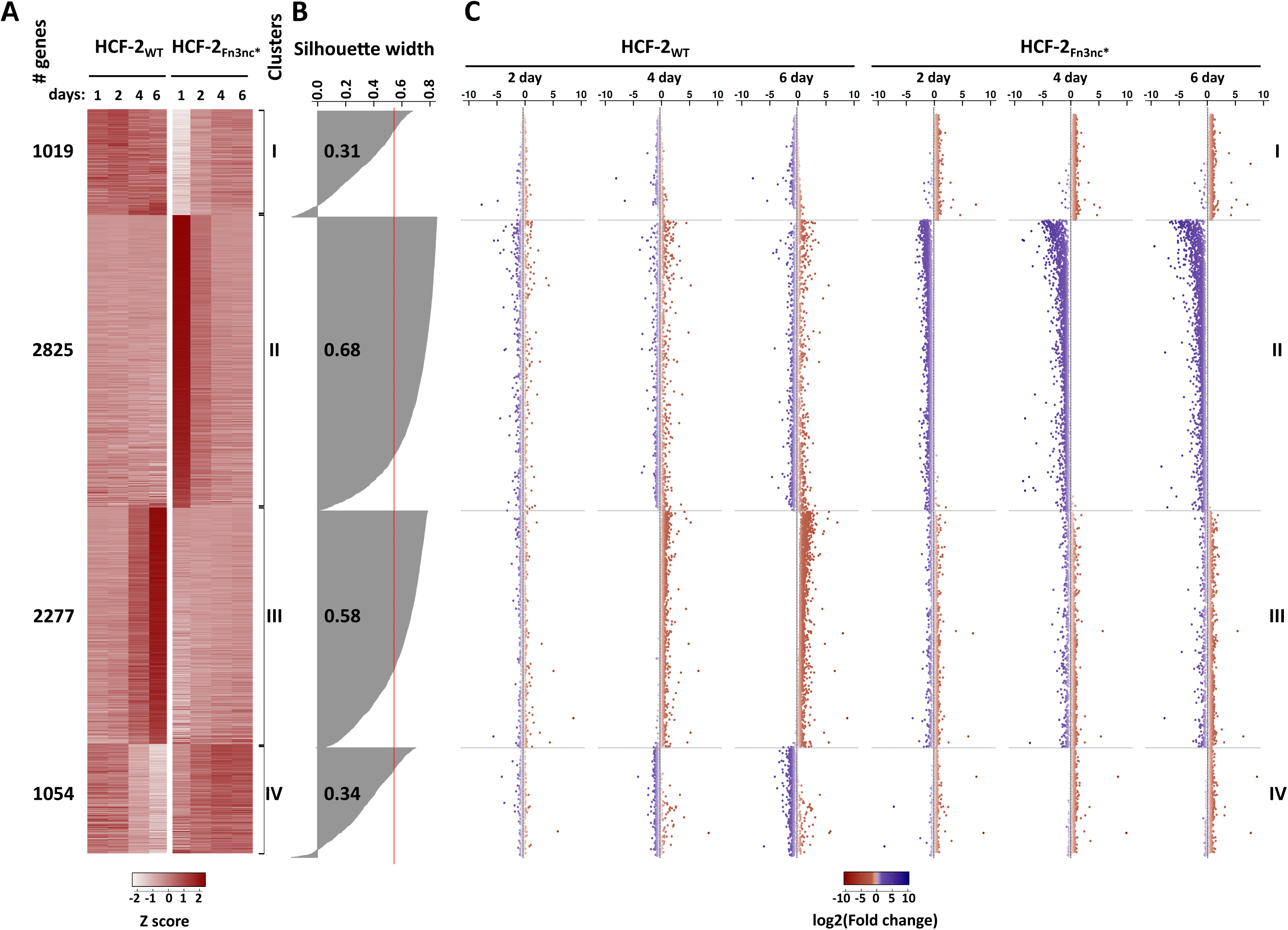
Induced HCF-2 synthesis leads to up-regulation of development-associated genes and down-regulation of metabolic genes. ***(A)*** Normalized counts of 7175 differentially expressed genes (see text, and Materials and Methods) were used to group genes into four clusters (I-IV, right) by expression profiles with PAM algorithm. The scaled and centered mean of the two replicates of each timepoint/condition is shown in the heat-map, with relative abundance of transcript per gene (Z-score) color-coded as shown in color scale. Left, number of genes per cluster. ***(B)*** Silhouette widths of each PAM-generated gene cluster. Red line, average for all clusters Silhouette width; fractional numbers, average Silhouette width for each individual cluster. ***(C)*** Scatter plot of log2(Fold change) of transcript abundance at days 2, 4, and 6 with respect to day 1. Genes are shown in the same order as in the heat-map.

In general, clusters I and II represent genes with little change in the HCF-2_WT_ samples but instead with up-regulation (in cluster I) and down-regulation (in cluster II) in the HCF-2_Fn3nc*_ mutant sample. Therefore, we focused on clusters III and IV where there were changes in the HCF-2_WT_ sample.

Cluster III displays a progressive increase in gene expression in the HCF-2_WT_ samples from days 1 to 6 with no dominant pattern in the HCF-2_Fn3nc*_ mutant. Gene Ontology (GO) analysis (Supplementary Table S3) followed by REVIGO graphic representation of cluster III (Figure 8A) revealed an enrichment of terms linked to extracellular organization, morphogenesis, developmental processes, and cell differentiation.

**Figure 8.**
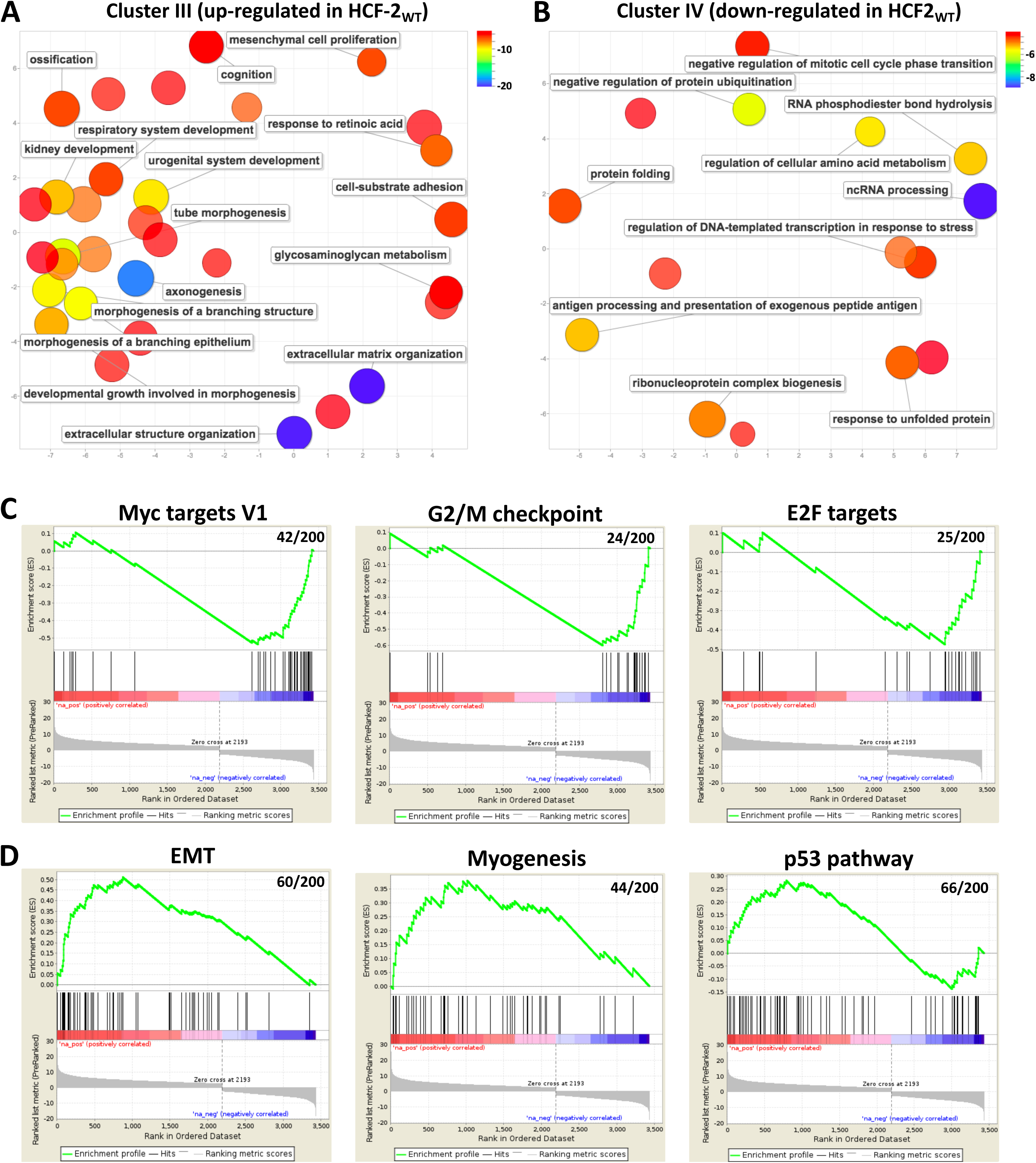
REVIGO display of GO terms associated with genes enriched in ***(A)*** cluster III (up-regulated in the HCF-2_WT_ sample) and ***(B)*** cluster IV (down-regulated in the HCF-2_WT_ sample). List of GO-terms generated by cluster Profiler (see Supplementary Table S3) and selected for a *p* value less than 10^−06^ for cluster III and less than 10^−05^ for cluster IV was analyzed using the REVIGO tool with an allowed similarity parameter of 0.7. GO-term bubbles are colored with respect to *p* values as indicated in the associated color scales, bubble size indicates the frequency of the GO term in the underlying GOA database. ***(C, D)*** Selected GSEA enrichment plots for Hallmark gene sets of differentially expressed genes of day 1 and day 6 of the HCF-2_WT_-samples for ***(C)*** down-regulated and ***(D)*** up-regulated pathways. Only the hallmarks most relevant to observed phenotype are shown (for more details of GSEA Hallmark sets see Supplementary Table S4 and Supplementary Figure S9).

In contrast, cluster IV largely represents genes down-regulated in the HCF-2_WT_ samples with some slightly increasing with the HCF-2_Fn3nc*_ mutant. Here, the top enriched GO terms were involved in metabolic processes including ncRNA processing, regulation of ubiquitination, and nucleotide and amino acid metabolism (Figure 8B, Supplementary Table S3).

Using Gene Set Enrichment Analysis (GSEA) with the set of 50 Hallmark pathways from the gene set collections described in (49), we analyzed the genes defined as differentially expressed by DESeq2 between day 1 and day 6 in either the HCF-2_WT_ (3440 genes) or HCF-2_Fn3nc*_ (4068 genes) samples. Genes were ranked by t-statistic, where a positive t-statistic indicates that the gene is more expressed in day 6 compared to day 1 (Figure 8C and 8D, Supplementary Figure S9, Supplementary Table S4).

Consistent with the decreased proliferation and increased G2/M defects of the HCF-2_WT_ cells (Figure 6), MYC and E2F targets, G2/M checkpoints, and DNA repair hallmarks were in general down-regulated (Figure 8C; Supplementary Figure S9A, Supplementary Table S4). Furthermore, in line with the reduced metabolism of less proliferative cells, mTORC1 signaling was also found down-regulated in HCF-2_WT_ - expressing cells (Supplementary Figure S9). Up-regulated GSEA Hallmark sets, in addition to Interferon α and γ response consistent with the study (15) (Supplementary Table S4), included prominent development related ones (e.g., epithelial-mesenchymal transition (EMT), myogenesis, angiogenesis, Hedgehog and Notch signaling; Figure 8D; Supplementary Figure S9B and S9C). The p53 pathway hallmark was also up-regulated in HCF-2_WT_, which is in line with the disturbance in cell-cycle progression.

In summary, elevated HCF-2 levels appear to result in inhibition of cell proliferation and metabolism, and activation of differentiation-related gene expression programs such as EMT.

## DISCUSSION

HCF-1 and HCF-2 represent related proteins encoded by a pair of paralogous genes, in humans called *HCFC1* and *HCFC2*; they most likely resulted from gene duplication around the time of divergence of the jawed gnathostome vertebrate lineage (Figure 1). Paralogs can often evolve to be expressed in the same cell but be responsible for different, even opposing functions, such as in the cases of *MYC* and *MXD1* (50), *E2F1* and *E2F4* (51), and *ELK1* and *ETS1* (51). Apparently, *HCFC1* and *HCFC2* are no exception, as here we demonstrate that HCF-2, like HCF-1, is broadly present, but in contrast to HCF-1, has acquired functions involved in inhibition of cell proliferation; it also can activate developmental gene expression programs. More striking, the subnuclear localization of HCF-1 and HCF-2 differ greatly, with the latter possessing strong nucleolar localization properties. These differing attributes are summarized in Figure 9A.

**Figure 9.**
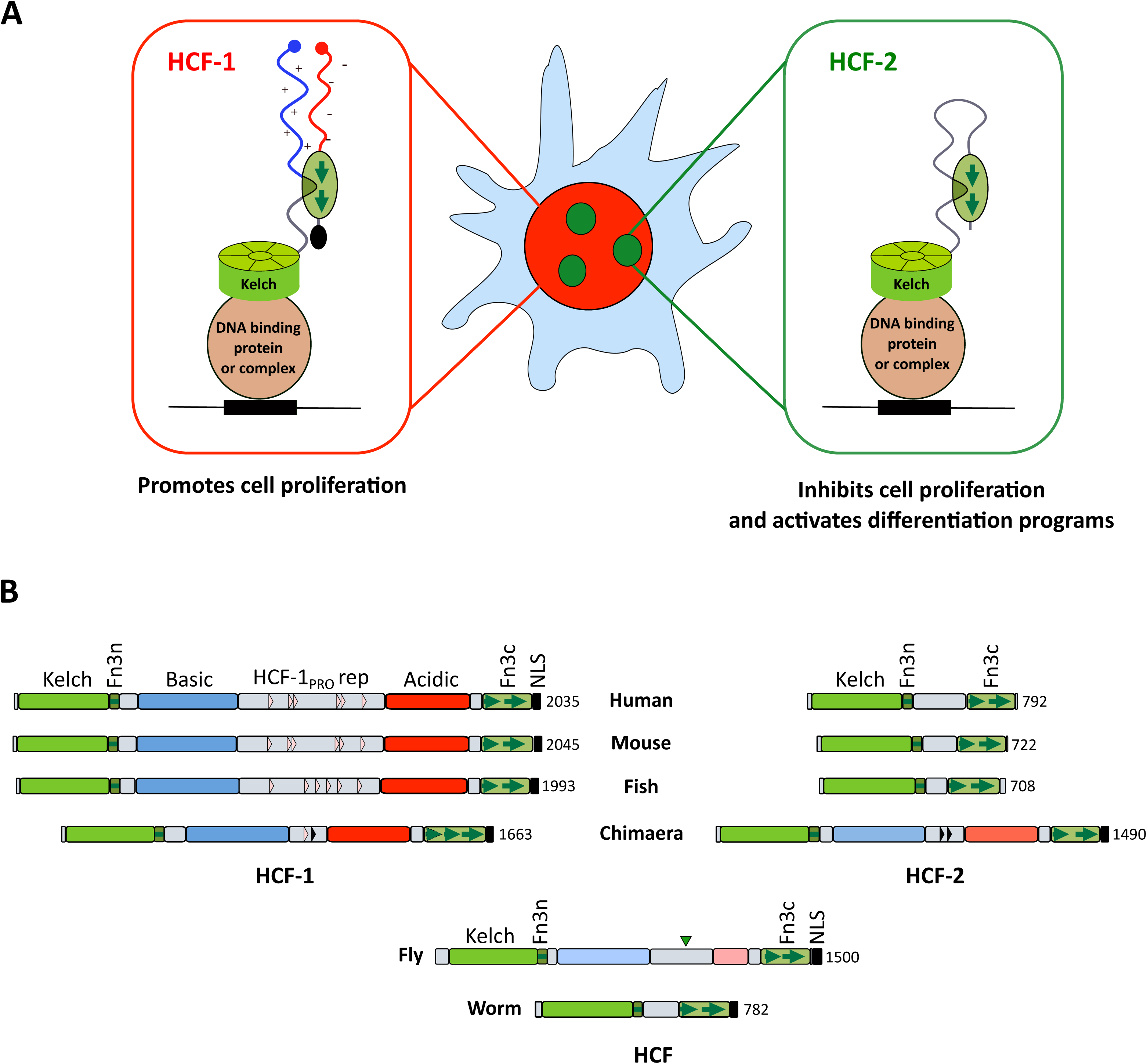
(***A***) Illustration of the different properties of human HCF-1 and HCF-2 proteins (see text for details). (***B***) Structural representation of HCF protein structures in worm, fly, chimaera, fish, mouse and human. For the domain descriptions see text and Figure 1A legend. Inverted arrow head represents a Taspase 1 cleavage site in fly HCF; black triangles in chimaera HCF-1 (one) and HCF-2 (two) indicate degenerate HCF-1_PRO_ repeats. The intensity of the blue and red colors labeling the Basic and Acidic regions indicates relative basic and acidic residue abundance: human, mouse, fish HCF-1 > chimaera HCF-1 and HCF-2 > *Drosophila* HCF.

Figure 9B shows a comparison of the structures of HCF-1 and HCF-2 in selected vertebrates with those of the single HCF proteins of two invertebrates: worm and fly. All of the HCF proteins retain certain characteristics: a Kelch domain, and Fn3n and Fn3c elements separated by a non-conserved element. During invertebrate evolution, HCF proteins evolved complexity between the Fn3n and Fn3c elements (e.g., Basic and Acidic regions flanking a Taspase 1 cleavage signal in *D. melanogaster*). Vertebrate HCF-1 retained this complexity, even acquiring the OGT HCF-1_PRO_-repeat cleavage sites, whereas in bony vertebrates HCF-2 evolved “simplicity,” keeping the Fn3n and Fn3c linker short and with no evident domain. Indeed, apparently, this inter Fn3n and Fn3c region lacks critical functions as rodents and fish miss a significant part of it.

Analysis of cartilaginous jawed vertebrates such as chimaera and shark reveals an illuminating intermediate form of HCF-2 lying between human HCF-1 and HCF-2 in structure as shown for chimaera in Figure 9B. Their Kelch domains segregate into HCF-1- and HCF-2-like (Figure 1B) and yet both chimaera (and shark) HCF proteins retain HCF-1-like Basic and Acidic regions, and NLS. Furthermore, both chimaera proteins possess two HCF-1_PRO_ repeat-like sequences (shown in Supplemental Figure S10); of which at least one of these — HCF-1_PRO_ repeat #1 in the HCF-1-like protein — is certainly cleavable by OGT. Thus, apparently before gene duplication, an ancestral HCF protein acquired the OGT HCF-1_PRO_ repeat cleavage signal and subsequently the Basic, Acidic, HCF-1_PRO_ repeat and NLS elements were lost in HCF-2 after gene duplication. Meanwhile, HCF-2 acquired nucleolar localization properties mediated by the conserved Fn3c element but also apparently dependent on the loss of the NLS, a shared feature of bony vertebrate HCF-2 proteins. Thus, HCF-2 acquisition of nucleolar localization properties via Fn3c evolution and NLS loss were likely important early proximate steps in HCF-2 evolution.

The nature and role of HCF-2 nucleolar localization are less clear. The presence of HCF-2 in the FC of the nucleolus where the Pol I subunit RPA194 resides and Pol I transcription is active, suggests a possible role in rRNA synthesis or early processing and, therefore, in ribosome synthesis generally. It is also possible, however, that HCF-2 has no direct role in nucleolar function, but rather that its nucleolar localization represents nucleolar “detention” (52), a mechanism by which specific proteins are sequestered from the cytoplasm or nucleus during cellular stress or certain cell metabolic states (e.g., Mdm2, VHL, POLD1). Nucleolar immobilization of such proteins is often mediated by their interaction with the transcript of the large spacer region that separates rRNA genes (53), consistent with HCF-2/RPA194 co-localization.

The similarity of the HCF-1 and HCF-2 Kelch domains — for example, both can interact with the HSV VP16 protein and under appropriate conditions (e.g. with VP16ΔC) form a transcriptional protein DNA complex (Figure 2) — suggests that, if ever released from the nucleolus, HCF-2 could either compete with HCF-1 for HCF-1-effector proteins or co-opt them, thus explaining the similar and respective loss-of-function and gain-of function HCF-1 and HCF-2 phenotypes. Consistent with this hypothesis, HCF-1 effectors (e.g., Set1/Ash2, Bap1, THAP11) can be found among HCF-2-associated proteins by MS analysis (Supplementary Table S1). Additionally or alternatively, HCF-2 may have acquired its own HCF-1-independent transcriptional regulatory roles and target other effector proteins, such as IRF-1 and IRF-2 as shown by Sun et al. (15).

Transcriptome analysis after induced HCF-2_WT_ synthesis revealed decreased expression of genes involved in metabolic processes (Figure 8B) and increased expression of genes related to differentiation and morphogenesis (Figure 8A), both being activities lost with the HCF-2_Fn3nc*_ mutant. These activities may result from the non-nucleolar — albeit nuclear — HCF-2 observed upon induced HCF-2_WT_ synthesis (Supplementary Figure S5A); such HCF-2 could interfere with or circumvent nuclear HCF-1 activities. Consistent with this hypothesis, many of the GO terms associated with down-regulation following induced HCF-2 synthesis were also found down-regulated in HeLa cells upon HCF-1 depletion (54), a result consistent with opposing cellular roles of HCF-1 and HCF-2. Such opposing roles may result from lost elements in HCF-2. For example, the human HCF-1 Basic region — missing in HCF-2 — is important for the ability of HCF-1 to promote cell proliferation (36, 55), and its Acidic region — also missing in HCF-2 — is important for its ability to promote proper M-phase progression (56) and for transcriptional activation (57). Thus, their disappearance in HCF-2 may be one reason that induced HCF-2 synthesis leads to retarded cell proliferation and elevated M-phase defects.

The activation of differentiation-related programs by HCF-2 shown here suggests novel activities separate from HCF-1. Interestingly, an analysis of publicly available RNA-seq databases demonstrates that in fish, frog and mouse embryos there are peaks of *HCFC2* gene expression during early embryogenesis (see Supplemental Figure S11). Indeed, a peak of *HCFC2* expression around gastrulation slightly precedes EMT, a gene expression program activated by HCF-2 in our studies.

Thus, we propose that HCF-2 is involved in activation of differentiation and morphogenesis gene expression programs, and in parallel in inhibition of cellular growth and metabolism. We believe that nucleolar localization of HCF-2 could play a role in regulation of these HCF-2 functions.

## DATA AVAILABILITY

The RNA-seq dataset generated in the study have been deposited in NCBI Gene Expression Omnibus (GEO) under accession number GSE128076. Token for reviewers: gvixyoeivpmvdap.

## Supporting information

Description

## ACKNOWLEDGMENTS

We thank Patrice Waridel, Manfredo Quadroni and the University of Lausanne Protein Analysis Facility for mass spectrometry analysis; Hannes Richter, Johann Weber and the Lausanne Genomic Technology facility for the high throughput RNA-sequencing; the Cellular Imaging Facility for assistance with the confocal microscope; Philippe Lhôte for aide with cell culture; Fabienne Lammers for the guidance with EMSA; Nouria Hernandez and Maria Cristina Gambetta for comments on the manuscript.

