## Supplemental Figure 1 for "HCF-2 inhibits cell proliferation and activates differentiation-gene expression programs"

### Figure S1

A

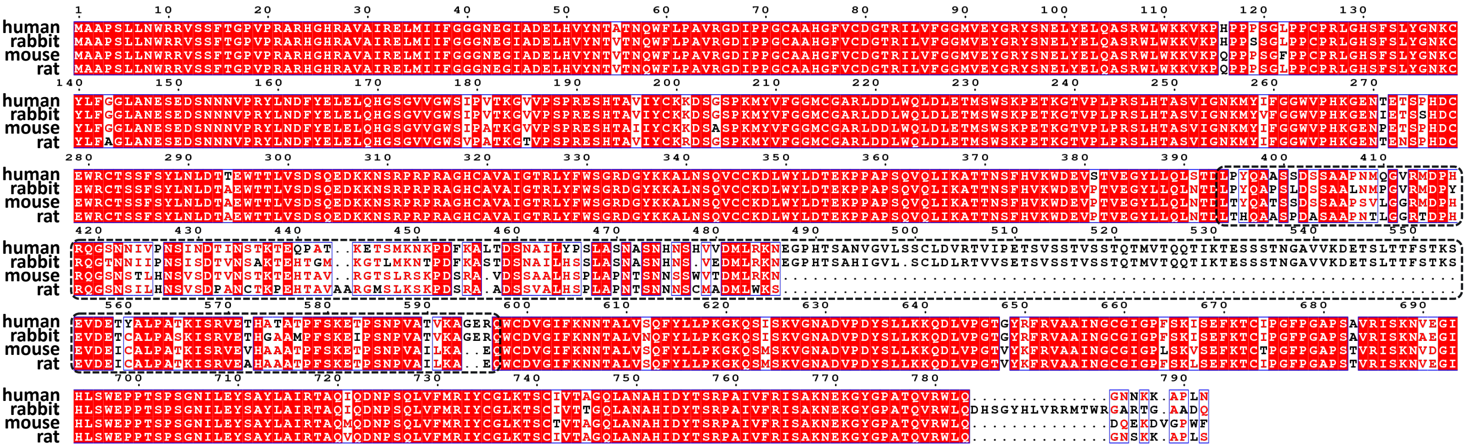


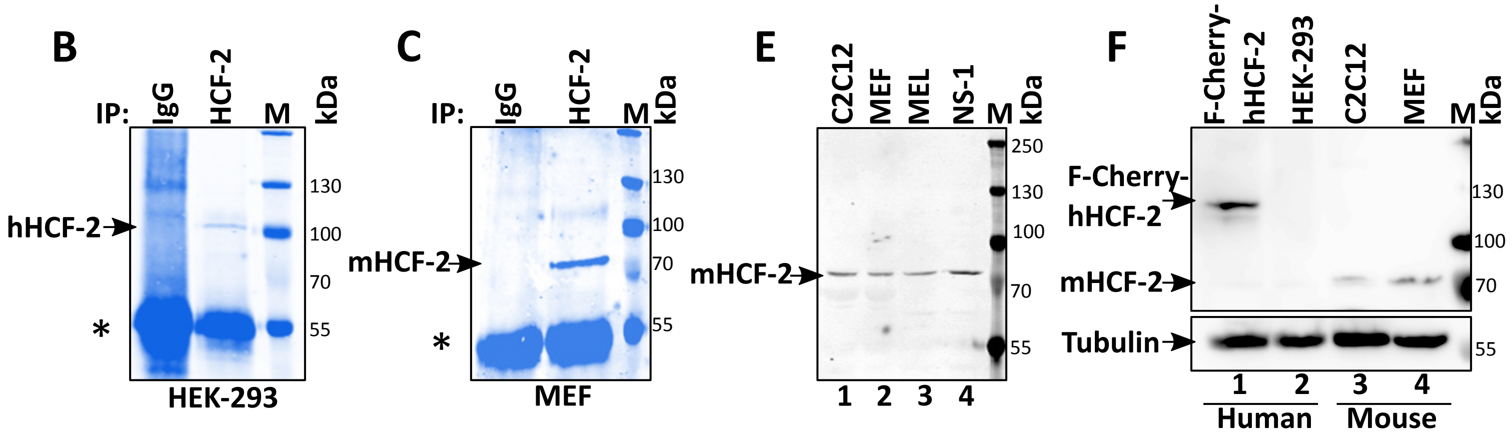


D

**IP from HEK-293:**

| | Identified proteins | Gene symbol | Molecular weight | Spectral counts vs IgG |
|---|---|---|---|---|
| 1 | Host cell factor 2 | HCFC2_HUMAN | 87 kDa | 121 |
| 2 | BTB/POZ domain containing KCTD3 | KCTD3_HUMAN | 89 kDa | 103 |
| 3 | Nucleolar RNA helicase 2 | DDX21_HUMAN | 87 kDa | 100 |
| 4 | Centromere protein F | CENPF_HUMAN | 368 kDa | 89 |
| 5 | RNA helicase DHX15 | DHX15_HUMAN | 91 kDa | 80 |

**IP from MEF:**

| | Identified proteins | Gene symbol | Molecular weight | Spectral counts vs IgG |
|---|---|---|---|---|
| 1 | Host cell factor 2 | G5E837_MOUSE | 79 kDa | 59 |
| 2 | 78 kDa glucose-regulated protein | GRP78_MOUSE | 72 kDa | 17 |
| 3 | Prelamin-A | LMNA_MOUSE | 74 kDa | 12 |
| 4 | RNA-binding protein 14 | RBM14_MOUSE | 69 kDa | 6 |
| 5 | Stress-70 protein, mitochondrial | GRP75_MOUSE | 73 kDa | 6 |
