## Supplementary figures and images for "HCF-2 inhibits cell proliferation and activates differentiation-gene expression programs"

### Supplemental Figure 2

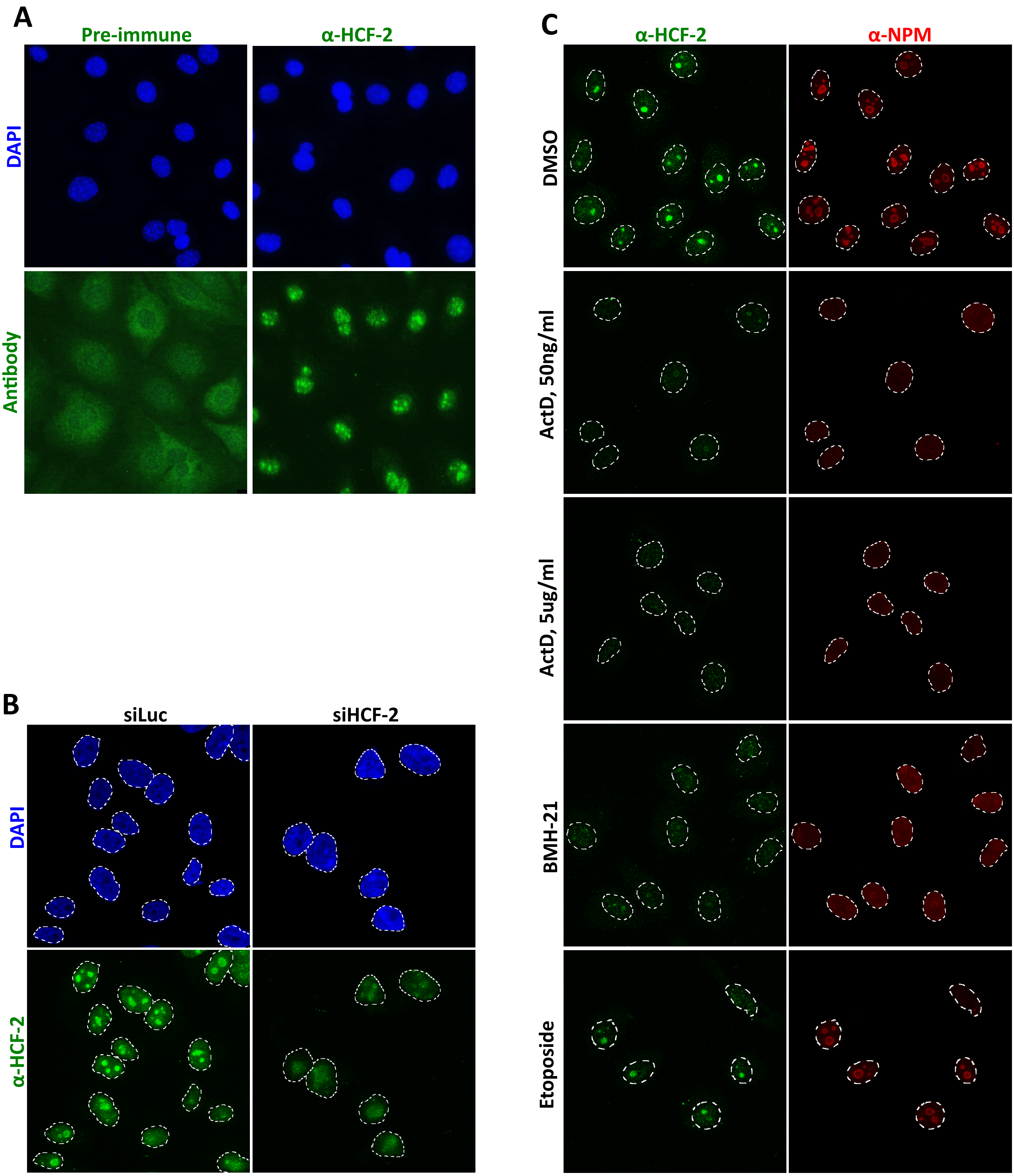

### Supplemental Figure 3

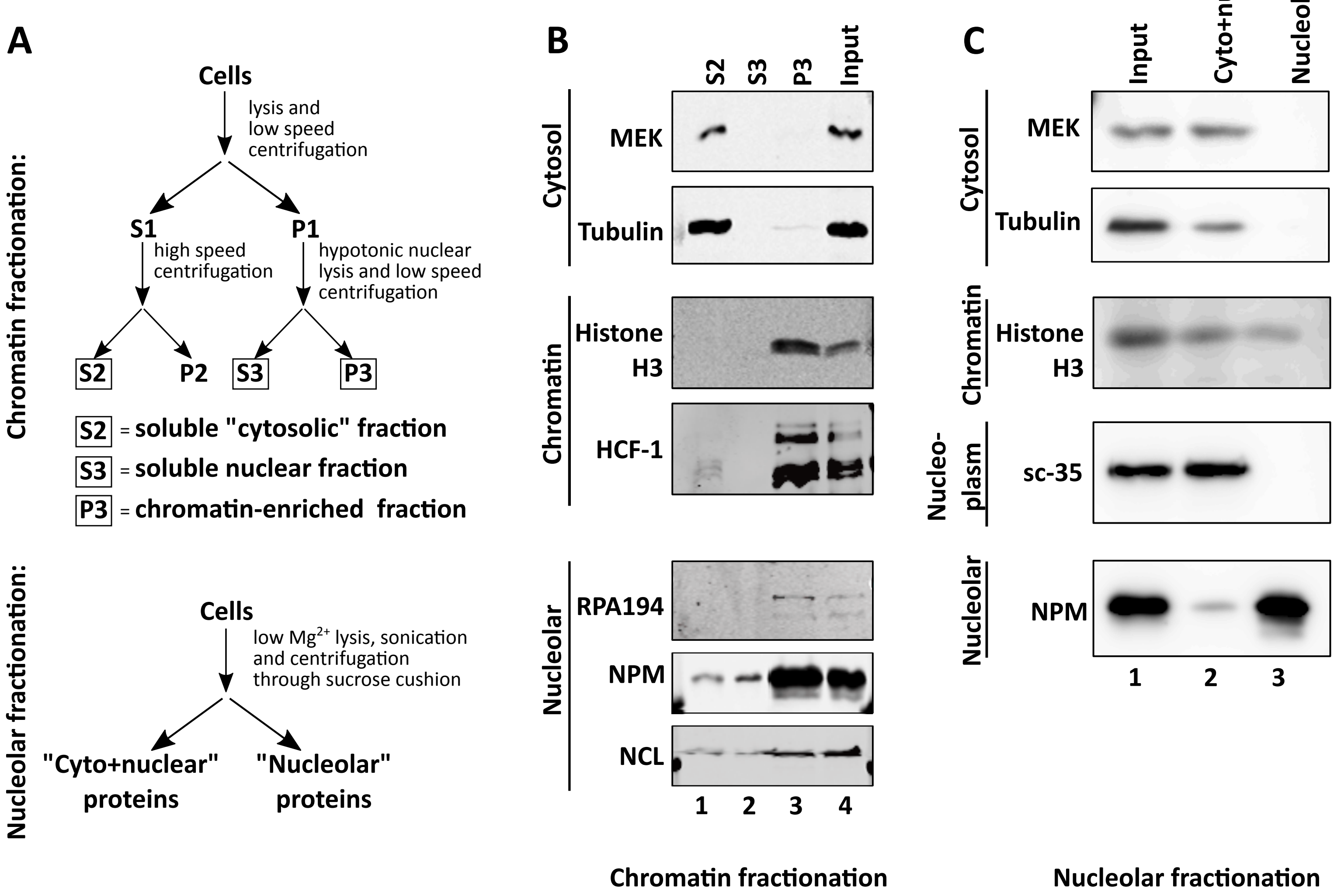

### Supplemental Figure 4

#### Figure S4

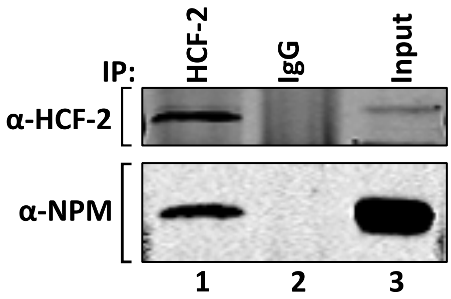

### Supplemental Figure 6

#### Figure S6

A

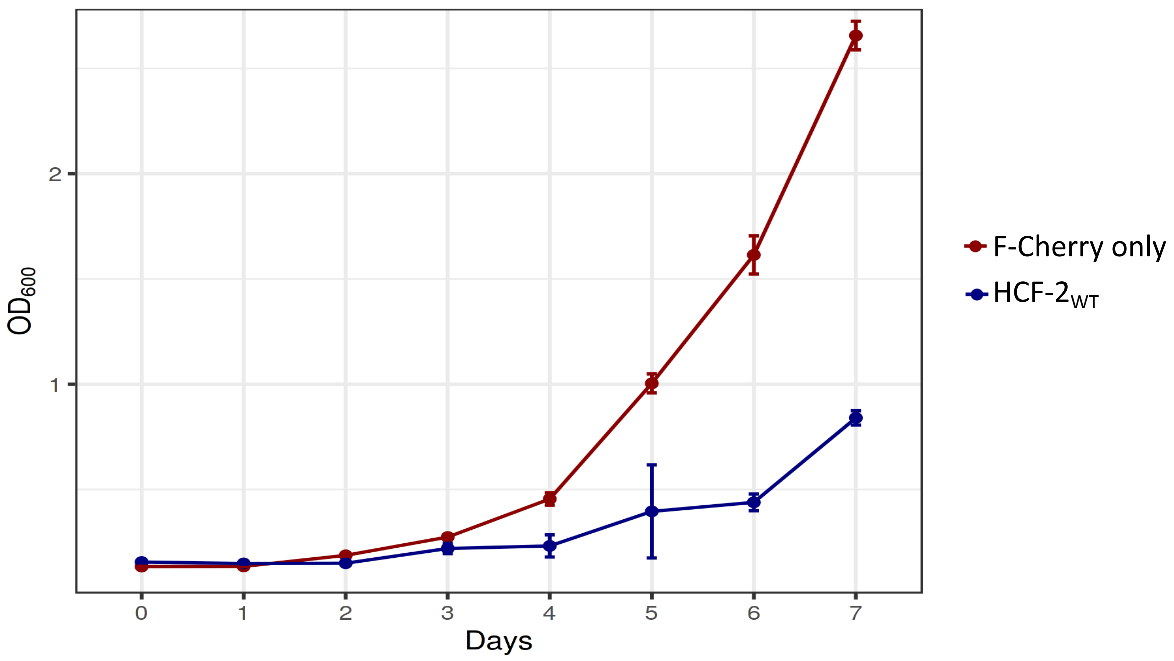


B

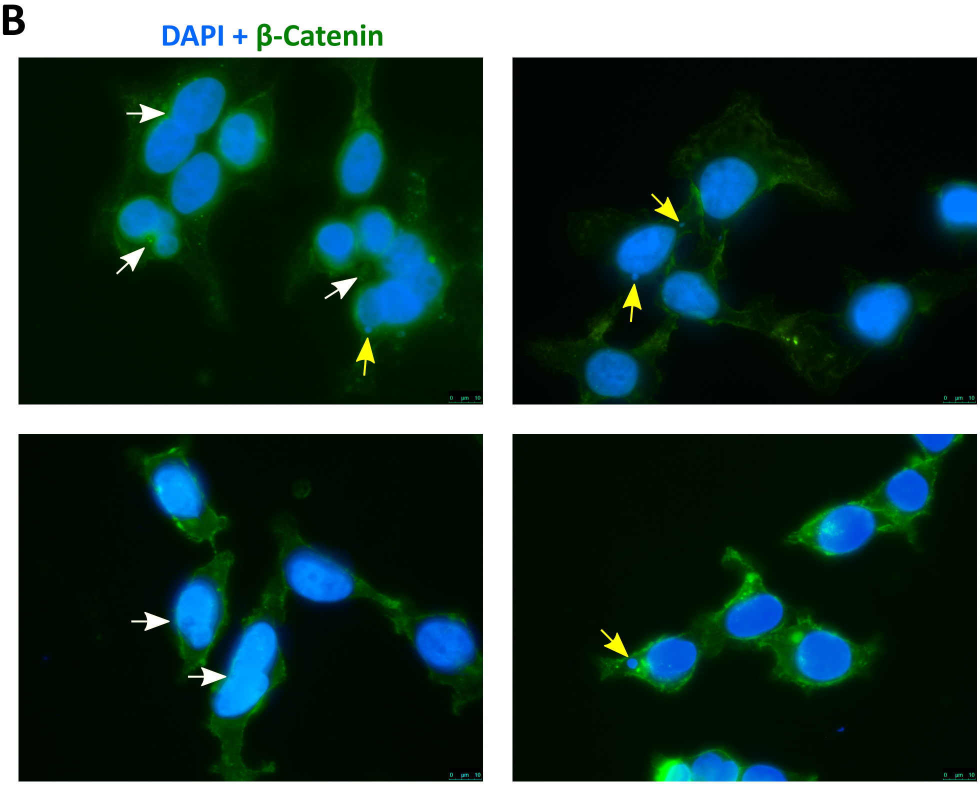

### Supplemental Figure 7

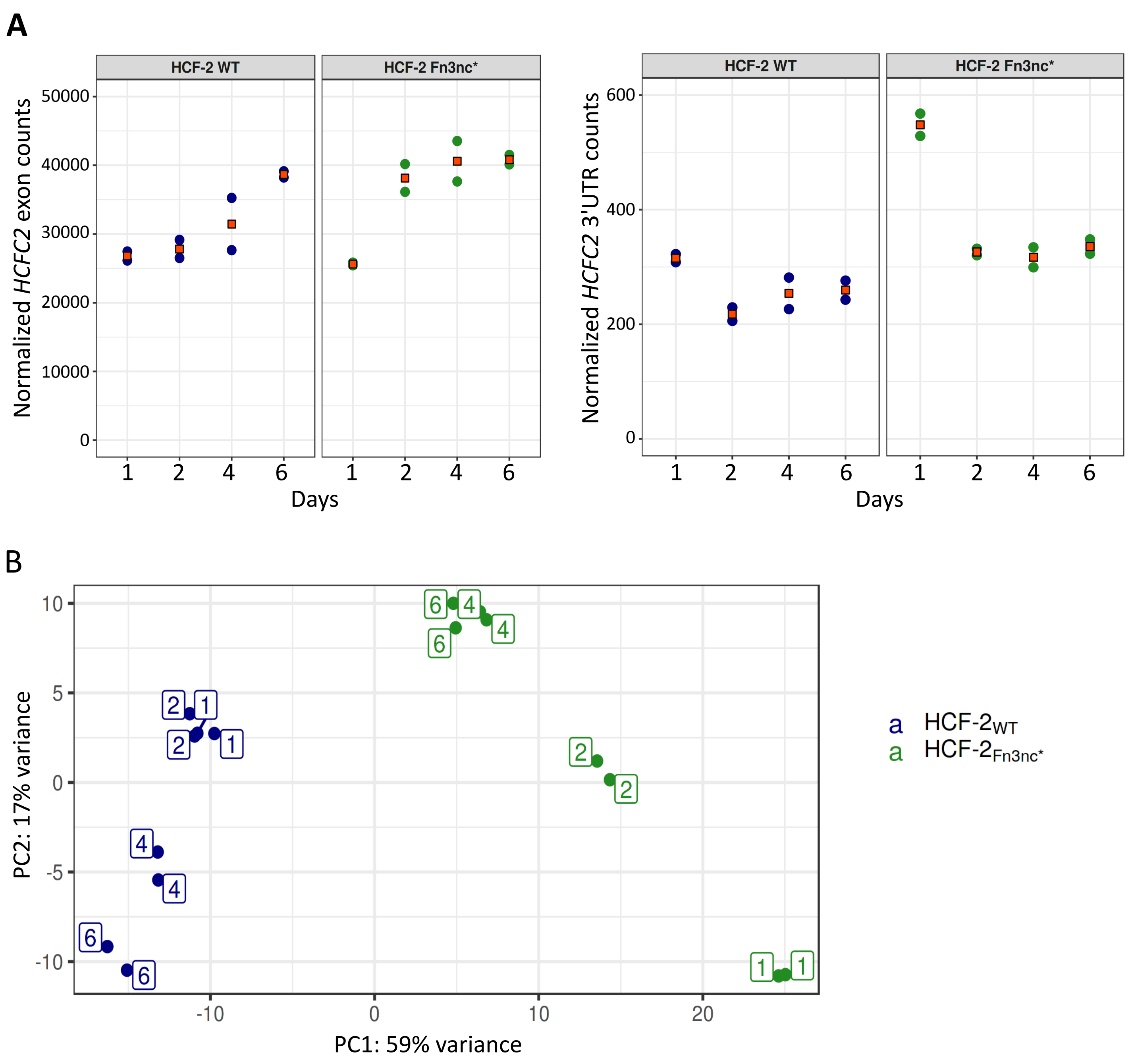

### Supplemental Figure 8

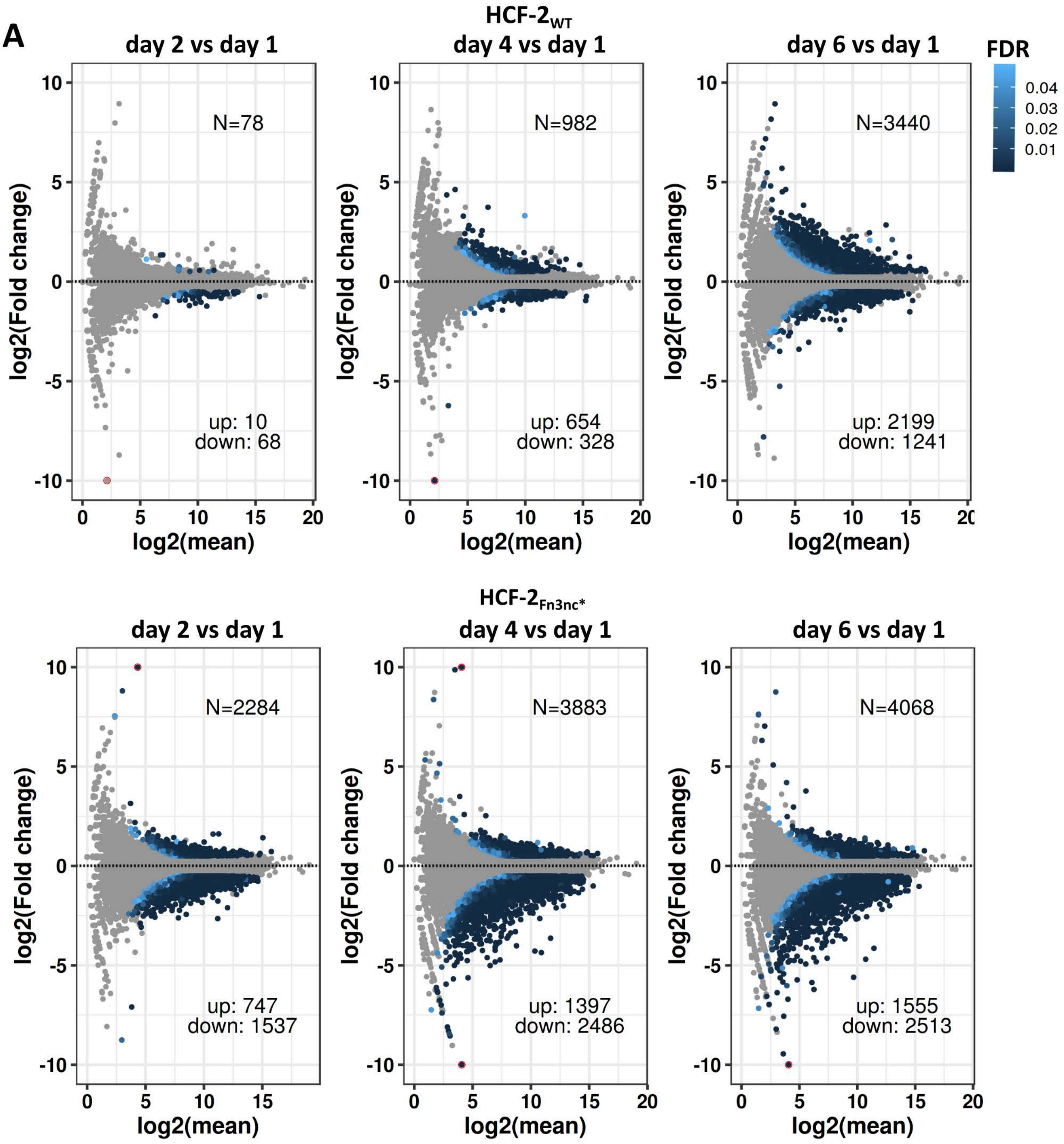

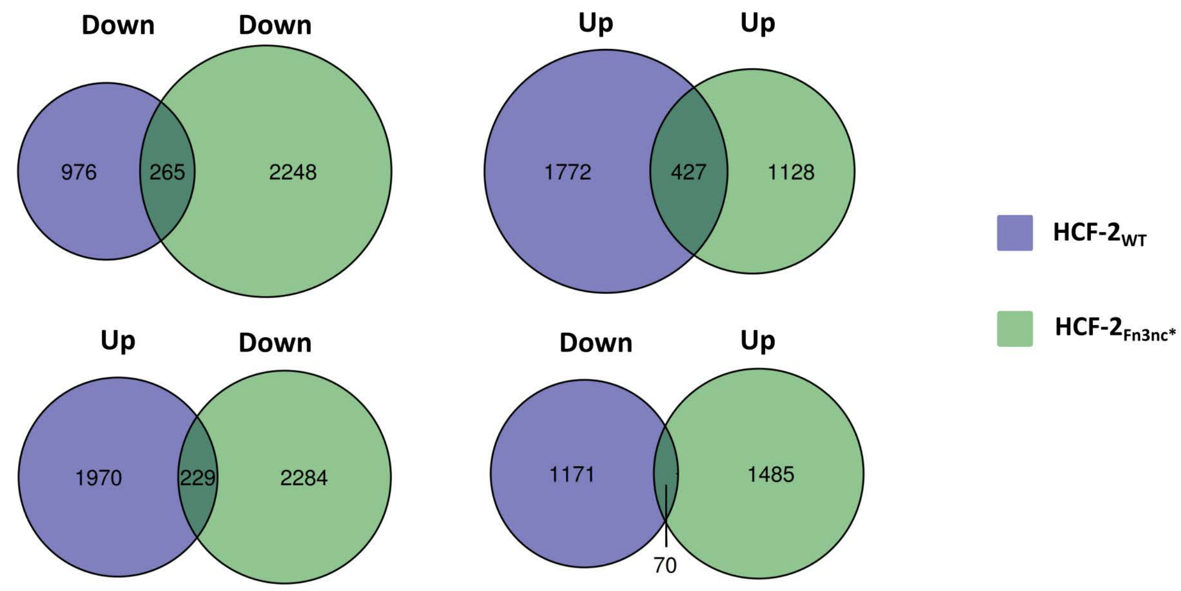

### Supplemental Figure 9

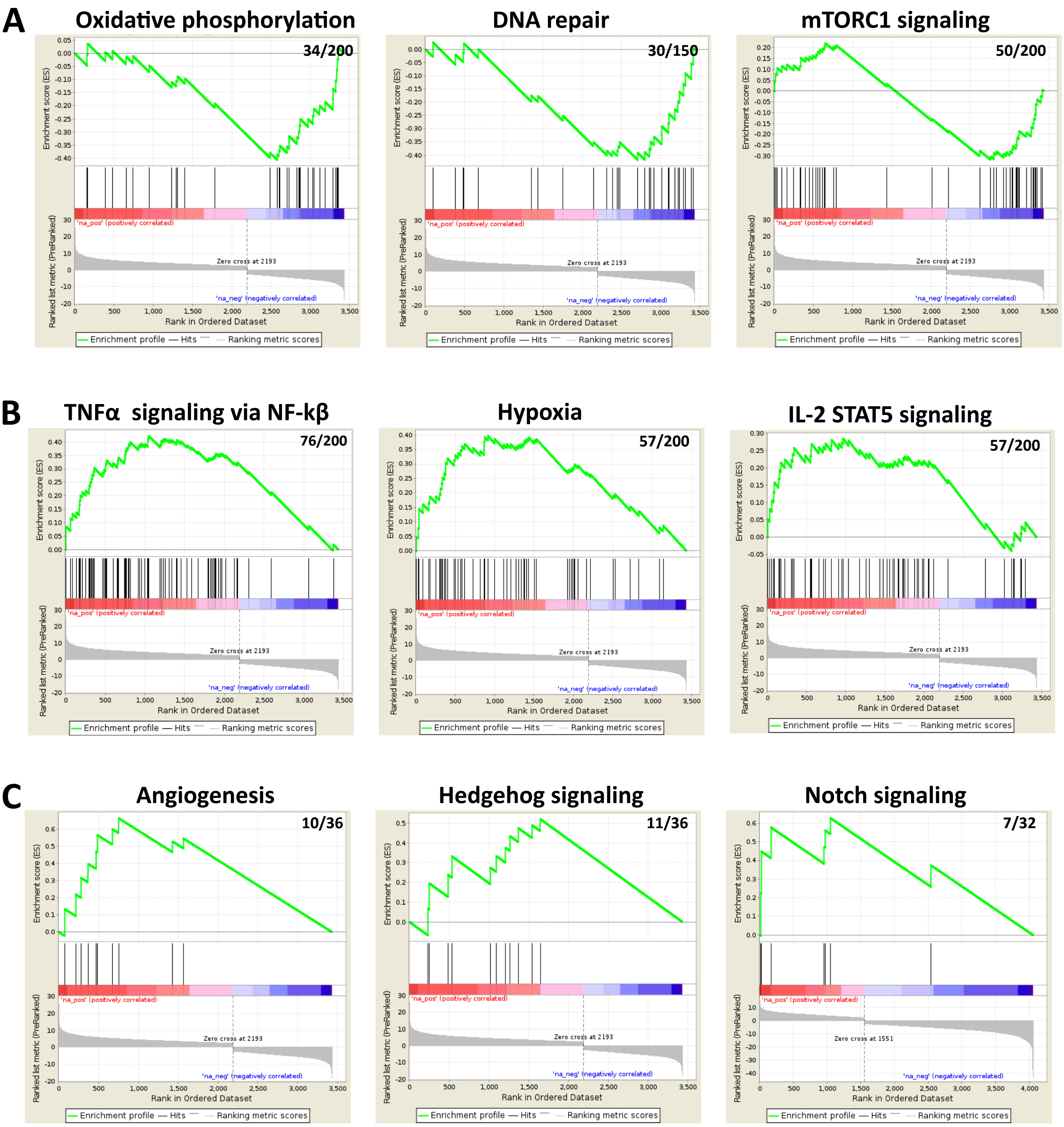

### Supplemental Figure 10

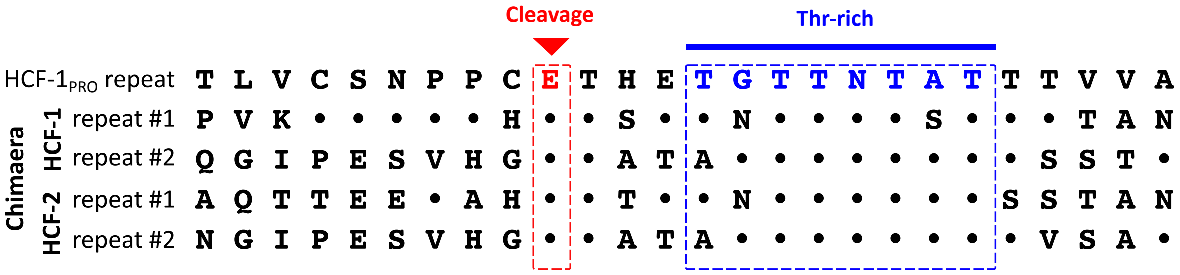

### Supplemental Figure 11

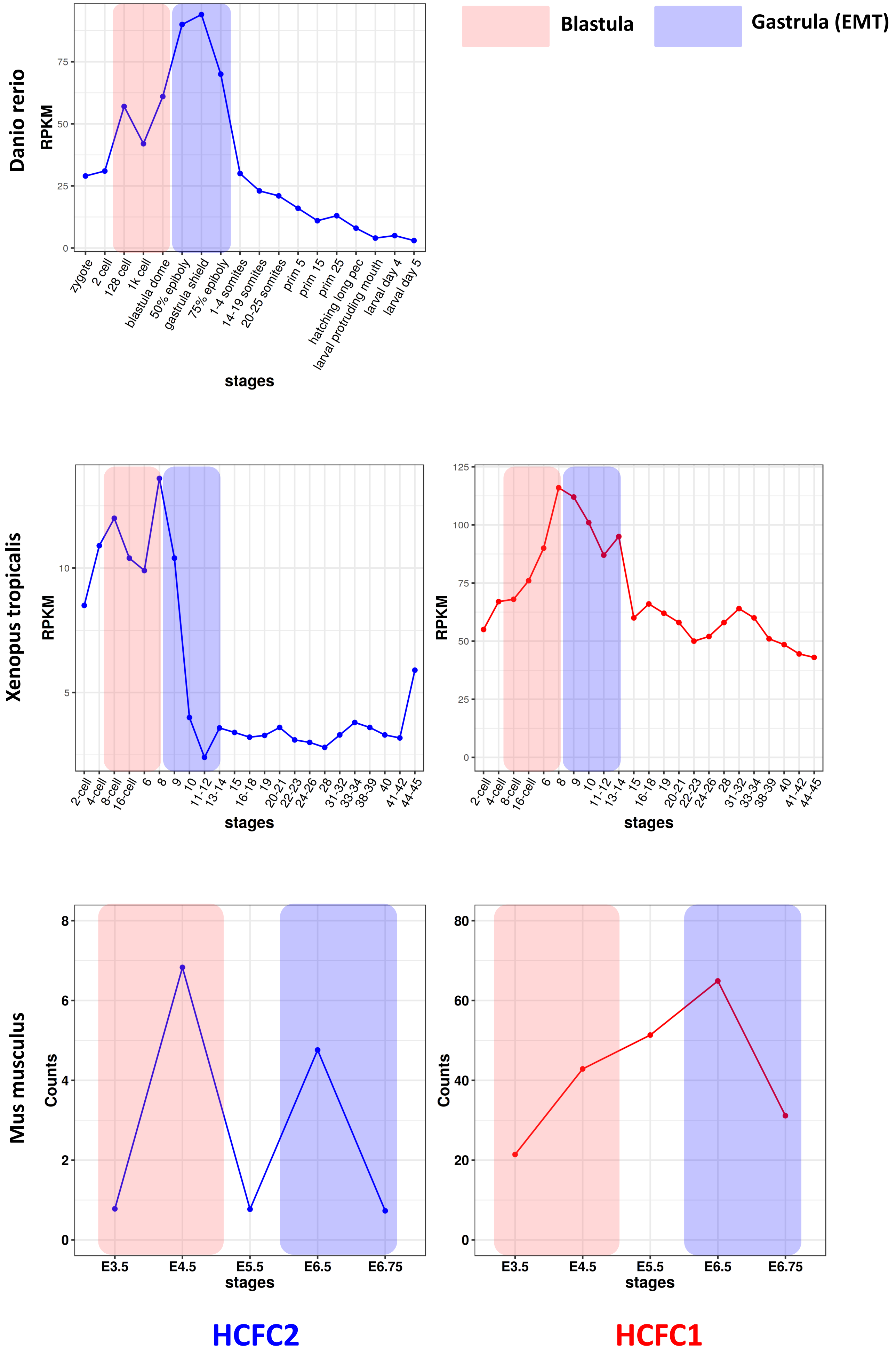
