## Supplemental Figure 5 for "HCF-2 inhibits cell proliferation and activates differentiation-gene expression programs"

### Figure S5

**A**

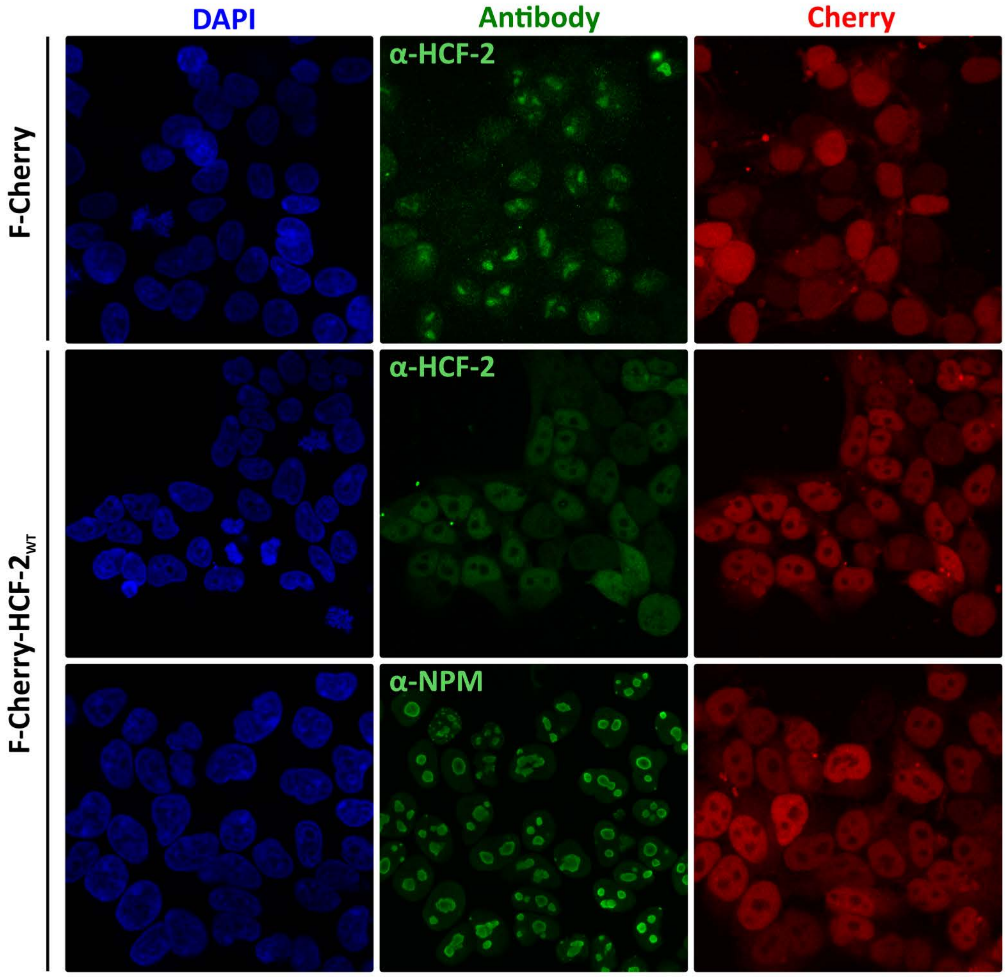


**B**

```
HCF-2  TEKPPAPSQVQLIKATTNSFHVKWDEVSTVEGYLLQL                          Fn3-1n
       TEKPP P++VQL++A TNS  V W  V+T + YLLQL
HCF-1  TEKPPPPARVQLVRANTNSLEVSWGAVATADSYLLQL

HCF-2  WCDVGICGNNTALVSQFYLLPKGKQSISKVGNADVPDYSLLKKQDLVPGTGYRFRVAAIN   Fn3-1c
       W DVG+      +V+  Y LP       S       VPDY+ LKKQ+L PGT Y+FRVA IN
HCF-1  WFDVGVIKGTNVMVTH-YFLPPDDAVPSDDDLGTVPDYNQLKKQELQPGTAYKFRVAGIN

HCF-2  GCGIGPFSKISEFKTCIPGFPGAPSAVRISKNVEGIHLSWEPPTSPSGNILEYSAYLAIR   Loop
        CG GPFS+IS FKTC+PGFPGAP A++ISK+ +G HL+WEPP+  SG I+EYS YLAI+
HCF-1  ACGRGPFSEISAFKTCLPGFPGAPCAIKISKSPDGAHLTWEPPSVTSGKIIEYSVYLAIQ

HCF-2  TAQI-----QDNPSQLVFMRIYCGLKTSCIVTAGQLANAHIDYTSRPAIVFRISAKNEKG   Fn3-2
       ++Q         P+QL FMR+YCG   SC+V +   L+NAHIDYT++PAI+FRI+A+NEKG
HCF-1  SSQAGGELKSSTPAQLAFMRVYCGPSPSCLVQSSSLSNAHIDYTTKPAIIFRIAARNEKG

HCF-2  YGPATQVRWLQ
       YGPATQVRWLQ
HCF-1  YGPATQVRWLQ
```

**C**

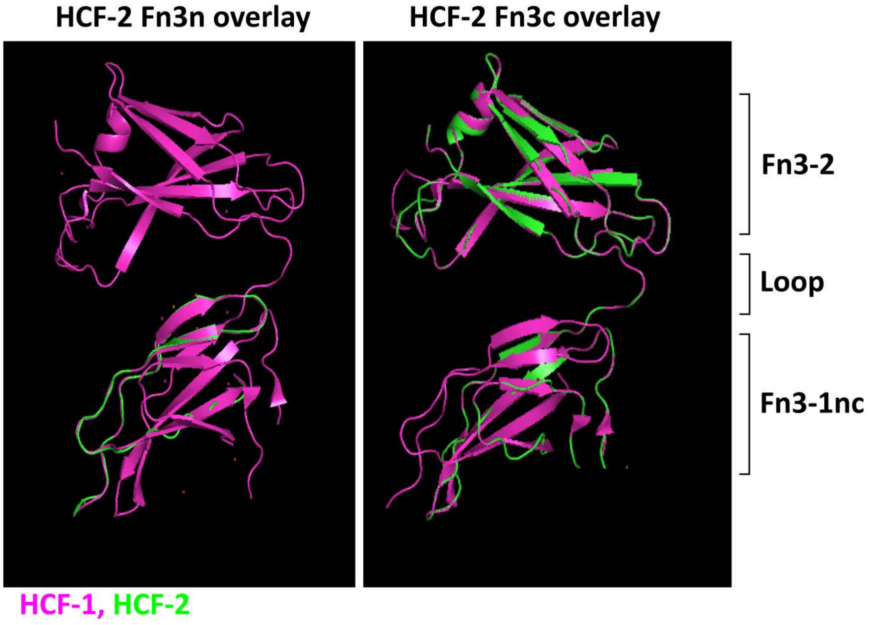
