## Supplementary material for "HCF-2 inhibits cell proliferation and activates differentiation-gene expression programs": Description

**SUPPLEMENTARY DATA**

**Supplementary Figures**

Figure S1. Related to Figure 1

Figure S2. Related to Figure 3

Figure S3. Related to Figure 3

Figure S4. Related to Figure 4

Figure S5. Related to Figure 5

Figure S6. Related to Figure 6

Figure S8. Related to Figure 7 and Figure S7

Figure S9. Related to Figure 8

Figure S10. Related to Figure 9 and Discussion

Figure S11. Related to Discussion

**Supplementary Tables (Separate files)**

Table S1. Related to Figure S1 and Figure S4

Table S2. Related to Figure 7, Figure S8

Table S3. Related to Figure 7, Figure 8

Table S4. Related to Figure 8, Figure S9

**SUPPLEMENTARY FIGURES**

**Figure S1.** ***(A)*** Alignment of human, rabbit, mouse, and rat HCF-2 protein sequences. The absence of amino acid sequences (488-555, in human HCF-2), encoding by exon 11 is rodent specific. The boxed sequence, representing the region of biggest sequence difference, depicts the mHCF-2 antigen used for rabbit immunization. ***(B, C)*** Coomassie-stained PAGE of immunoprecipitated (IP) hHCF-2 from HEK-293 cells (***B***) and mHCF-2 from MEF (***C***). Arrows identify the hHCF-2 and mHCF-2 bands of different size. IgG, non-immune serum. Asterisk, antibody heavy chain. ***(D)*** Results of MS of indicated bands in **B** and **C**. ***(E)*** Immunoblot of endogenous mHCF-2 from indicated mouse cell extracts (lanes 1-4) with affinity purified α-HCF-2 antibody. ***(F)*** Immunoblot of human and mouse HCF-2. Lane 1, Cherry-hHCF-2 fusion protein; lane 2, human HEK-293 cell extract; lanes 3 and 4, mouse C2C12 and MEF cell extracts, respectively. Tubulin, loading control.

**Figure S2. *(A)*** Immunostaining of MEF cells either with pre-immune serum (left bottom) or α-HCF-2 antibody (right bottom, green). Upper panels, nuclear DAPI staining (blue). ***(B)*** HEK-293 cells were treated with siRNA against luciferase (left) or HCF-2 (right) for 72 h and then stained with α-HCF-2 antibody (green, lower panels). The upper panels show cell nuclei stained with DAPI (blue). ***(C)*** HEK-293 cells were treated for 4 h with (from left to right) DMSO (vehicle), 50 ng/ml and 5 ug/ml actinomycin D (ActD), 1 uM BMH-21, and 25 uM etoposide. Cells were stained with α-NPM antibody (red, upper panels) for nucleolar integrity and α-HCF-2 (green, lower panels). Dashed line denotes the nuclear area.

**Figure S3.** Analysis of a small-scale chromatin and nucleolar fractionation procedures. **(*A*)** Schematics of chromatin (upper) and nucleolar (lower) fractionation procedures. **(*B*)** Chromatin fractionation of HEK-293 cells showing S2 (lane 1), S3 (lane 2), and P3 (lane 3) fractions and whole cell lysate (lane 4) stained with α-MEK and α-tubulin antibodies for cytosolic proteins; α-HCF-1 and α-histone 3 (H3) antibodies for chromatin-bound proteins; and α-RPA194, α-NPM and α-NCL antibodies for nucleolar proteins. **(*C*)** Nucleolar fractionation of HEK-293 cells showing whole cell lysate (lane 1); cytoplasmic and nuclear (lane 2), and nucleolar (lane 3) fractions. Protein markers are as in ***B***, except for sc-35 for the nucleoplasm.

**Figure S4.** Immunoblot of NPM in protein complexes immunoprecipitated from MEF extracts with α-HCF-2 antibody. Lane 1, IP with α-HCF-2 antibody; lane 2, IP with negative control IgG; lane 3, whole cell lysate.

**Figure S5. *(A)*** Co-immunostaining of HEK-293 cells expressing either F-Cherry (upper row) or F-Cherry-HCF-2_WT_ (middle and bottom rows) 24 h after induction with doxycycline with either α-HCF-2 or α-NPM antibodies (green). Left panels, nuclear staining with DAPI (blue), right panels, a signal from Cherry. ***(B)*** Alignment of amino acid sequences of Fn3n and Fn3c domains of hHCF-2 (upper line) with hHCF-1 (bottom line). The middle line represents identities (letter) and similarities (+) between the two sequences. The Fn3-1 sequences are boxed in light green and Fn3-2 sequences are boxed in dark green. The Fn3 repeats are linked by 6 amino acid loop (boxed in pink).  ***(C)*** PyMOL generated overlay of HCF-2 (green) Fn3n (left) and Fn3c (right) elements with HCF-1 (pink) Fn3nc structure (21).

**Figure S6. *(A)*** MTT assay of HEK-293 cells with either F-Cherry only or F-Cherry-HCF-2_WT_ during 7 days after addition of doxycycline. ***(B)*** F-Cherry-HCF-2_WT_ cells display a number of mitotic defects 72 h after doxycycline induction. Multinucleated cells (white arrows) and micronuclei (yellow arrows) are indicated.

**Figure S7. *(A)*** Normalized read counts mapped to (i) HCF-2-coding sequences (left) resulting from both endogenous and recombinant *HCFC2* sequences and (ii) *HCFC2* 3’UTR sequences (right) from endogenous *HCFC2* sequences only. **(B)** Plot of the two major PCA components of individual RNA-seq samples, dots’ labels indicate day after doxycycline addition.

**Figure S8. *(A)*** MA plots of read counts for genes in samples with F-Cherry-HCF-2_WT_ (upper row) and F-Cherry-HCF-2_Fn3nc*_ (lower row) of indicated day versus day 1. Mean of two replicates +1 pseudocount is shown. Genes with absolute value of log2FoldChange >10 were automatically rescaled to 10 are encircled with red. ***(B)*** Venn diagrams represent common and unique genes found differentially expressed in both F-Cherry-HCF-2_WT_ (blue) and F-Cherry-HCF-2_Fn3nc*_ (green) samples at day 6 with respect to day 1.

**Figure S9.** Selected GSEA enrichment plots for Hallmark gene sets with high absolute value of normalized enrichment scores (NES) based on Gene Set Enrichment Analysis of differentially expressed genes of day 1 and day 6 of the HCF-2_WT_-samples. Only the hallmarks most relevant to observed phenotype are shown (for full list of GSEA Hallmark sets see Supplementary Table S4). ***(A)*** Down-regulated and ***(B)*** up-regulated GSEA Hallmark sets. ***(C)*** Up-regulated Hallmark sets with fewer than 50 genes. Numbers in the right upper corner indicate ratio of genes found differentially expressed to total number of genes in hallmark gene set.

**Figure S10.** Comparison of a human HCF-1_PRO_-repeat sequence with two HCF-1_PRO_ repeat-like sequences in Chimaera HCF-1-like (1007-1031 and 1045-1070 aa) and HCF-2-like (920-945 and 959-984 aa) proteins. Cleavage site and Threonine-rich sequences are shown in red and blue respectively. Dot represents identity with human HCF-1_PRO_ repeat 1 sequence.

**Figure S11.** Analysis of public data sets for *HCFC2* expression (RPKM or counts) in fish ***(A)***, frog ***(B)*** and mouse ***(C)*** early embryonic stages (60-62). Respective developmental stages for each species are shown. Stages, corresponding to the formation of the blastula and the gastrula are boxed in pink and blue respectively.

**SUPPLEMENTARY TABLES**

**Table S1.** List of proteins identified by mass spectrometry of immunoprecipitated mHCF-2 complexes from MEF extracts. Ctrl, non-immune rabbit IgG

**Table S2.** List of genes detected in RNA-seq of HEK-293 cells with induced HCF-2_WT_ or HCF-2_Fn3nc*_ synthesis. Table S2 shows list of genes with FPKM > 1 which were kept for differential expression analysis done with DESeq2. Columns J and Z (entitled as “Average of expression") show averaged counts for samples with induced expression of HCF-2_WT_ and HCF-2_Fn3nc*_ respectively. Cluster identifiers produced by PAM (see Materials and Methods) are shown in column I and NA means that gene was not attributed to any of clusters, e.g. gene was stably expressed across all samples. lfcSE is standard error of log2Foldchange estimate; stat is t-statistic returned by Wald test from DESeq2 package. Genes’ genomic positions are shown in respect to GRCh37 (Hg19) genome version, gene annotations were obtained from www.gencodegenes.org.

**Table S3.** List of GO terms associated with genes in PAM clusters I–IV. GO terms with FDR < 0.05 are listed.

**Table S4.** GSEA report for differentially expressed gene sets in F-Cherry-HCF-2_WT_ and F-Cherry-HCF-2_Fn3nc*_ samples. Results of GSEA run on 50 hallmark pathways for sets of differentially expressed genes for HCF-2_WT_ or HCF-2_Fn3nc*_ between day 6 and day 1. ES is enrichment score, NES - normalized enrichment score. ORIGINAL SIZE column represents number of genes annotated to hallmark pathway, SIZE column gives number of differentially expressed genes per pathway. For details visit <https://software.broadinstitute.org/gsea/doc/GSEAUserGuideFrame.html>
